# Regulation of *Klebsiella pneumoniae* mucoidy by the bacterial tyrosine kinase Wzc

**DOI:** 10.1101/2022.06.05.494587

**Authors:** Saroj Khadka, Brooke Ring, Lindsey R. Krzeminski, Matthew Hathaway, Ryan S. Walker, Harry L.T. Mobley, Laura A. Mike

## Abstract

*Klebsiella pneumoniae* is a frequent nosocomial pathogen associated with urinary tract infections (UTI), pneumonia, and septicemia. Two lineages of *K. pneumoniae* have emerged that challenge the treatment of these infections. One lineage includes hypervirulent strains, causing invasive community-acquired infections, and the other encompasses carbapenem-resistant isolates. Non-mucoid, carbapenem-resistant isolates are most frequently isolated from UTI. Therefore, we hypothesized that environmental conditions may drive *K. pneumoniae* adaptation to the urinary tract. We found that two *rmp*-positive, hypervirulent strains and two *rmp*-negative clinical UTI isolates all suppressed mucoidy when cultured in urine compared to rich LB medium without altering capsular polysaccharide (CPS) abundance. We then performed a transposon screen to identify genetic factors through which urine suppresses mucoidy. Due to variations within transposon hits, we performed whole genome sequencing on our hits and found 100% association between secondary mutations in Wzc and increased mucoidy. Wzc is an inner membrane tyrosine kinase that regulates CPS assembly. When Wzc mutations are encoded chromosomally we observed both increased mucoidy and increased cell-free extracellular polysaccharide (EPS). However, when Wzc mutations are expressed *in trans* we observed increased mucoidy without impacting CPS or cell-free EPS abundance. Functionally, one periplasmic mutation increased Wzc autophosphorylation, while four active site mutations reduced Wzc autophosphorylation. These four active site mutations are also sufficient to increase mucoidy in one of the *rmp*-negative clinical UTI isolates without affecting EPS abundance. Combined, these data implicate mutation-driven modulation of Wzc activity as a global, *rmp*-independent mechanism for increasing *K. pneumoniae* mucoidy, without altering EPS uronic acid content. Wzc activity may represent the lynchpin that coordinates both capsule biosynthesis and mucoidy, explaining why these two phenotypes have been historically intertwined.

## INTRODUCTION

*Klebsiella pneumoniae* is a gram-negative pathogen responsible for approximately 10% of all nosocomial infections.^1^ Within the hospital setting, *Klebsiella* cause 23% of all urinary tract infections (UTIs), 14% of all surgical-site infections, 12% of all pneumonia cases, and 8% of all bloodstream infections.^1^ In the community setting, *K. pneumoniae* is the second most common cause of noncomplicated UTI after *Escherichia coli*.^2^ Isolates that pose a major threat to human health include carbapenem-resistant (CRKp) and hypervirulent (hvKp), which cause severe morbidities and high mortality.^3-5^ CRKp was first observed in 1996 and since then it has been the major driving force disseminating the carbapenem resistance cassette throughout the *Enterobacteriaceae*, complicating the treatment of gram-negative infections.^6,7^ Of all healthcare-associated infections, CRKp is most commonly isolated from UTI cases, challenging the therapeutic choices of healthcare-providers.^8-10^ In parallel, the incidence of hvKp is rising in both hospital and community settings.^11-14^ HvKp is associated with invasive infections, especially pyogenic liver abscesses that disseminate to the eyes and brain, a pathogenesis uncommon for gram-negative enteric bacteria.^3,4^ Features associated with hvKp include hypermucoviscosity, *rmpA* overexpression, and K1 or K2 capsular polysaccharide.^4,11,13,14^ However, some hvKp do not encode K1 or K2 capsule (CPS) and some K1 or K2-encapsulated strains are not hypervirulent.

The historical model of *K. pneumoniae* hypervirulence has been that overproduction of CPS boosts mucoidy and promotes resistance to host defenses. This is due to observations that elevated *rmpA* expression increases CPS production and mucoidy, and that loss of CPS production ablates mucoidy and virulence.^4^ However, recent studies have demonstrated that other factors contribute to mucoidy. An elegant dissection of the *rmp* locus identified its polycistronic structure consisting of the auto-regulator *rmpA* and two downstream genes, *rmpC* and *rmpD*, that distinctly impact CPS and mucoidy.^15^ An *rmpC* deletion strain retains mucoidy despite reduced CPS.^16^ Correspondingly, an *rmpD* deletion strain loses mucoidy and retains full CPS production.^17^ Overexpressing *rmpD* in acapsular strains does not restore mucoidy, supporting the model that mucoidy requires CPS biosynthetic components, plus other unknown biochemical factors.^17^ Our previous studies provide further support of this model, as we identified many genes that coordinately increase or decrease CPS and mucoidy.^18^ However, we also identified some metabolic genes that only affect CPS or mucoidy.^18^ Moreover, we identified no genes that support mucoidy in acapsular strains, further supporting that mucoidy requires CPS biosynthesis.

These previous studies were performed in the ATCC 43816 derivatives, KPPR1. This strain is a model hypervirulent and mucoid strain that produces a K2 CPS, comprised of glucose (Glc), mannose (Man), and glucuronic acid (GlcA). *K. pneumoniae* employ the Wzy-dependent pathway to synthesize their Group 1 CPS.^19^ Undecaprenyl diphosphate-linked CPS-subunits are assembled on the cytoplasmic side of the inner membrane by glycosyltransferases then flipped to the periplasm by Wzx. There, Wzy polymerizes the growing capsular polysaccharide chain. Continued polymerization requires coordinated Wzc tyrosine autokinase and Wzb phosphatase activity.^20^ Wzc is an integral inner membrane protein that forms a trans-periplasmic complex with the CPS export pore, Wza, in the outer membrane. Wzc phosphocycling resulting in octamer association (unphosphorylated state) and dissociation (phosphorylated state) coordinates Wza extrusion of the growing polysaccharide chain.^21^ Wzi then anchors the CPS to the cell surface by an unknown mechanism.^22,23^ Previous work identified CRKp isolates with increased mucoidy and CPS production in clinical bloodstream isolates; along with non-mucoid and acapsular strains in clinical UTI isolates.^24^ None of the isolates encoded the *rmp* locus, distinct from KPPR1. Whole genome sequencing revealed that point mutations in *wzc* increased CPS production and mucoidy, which promoted phagocytosis resistance, enhanced dissemination, and increased mortality in animal models. Conversely, disruptions in *wbaP* ablated CPS production and mucoidy, which improves epithelial cell invasion, biofilm formation and persistence in the urinary tract. These data demonstrate that mutation of Wzc is an *rmp-*independent pathway that can enhance mucoidy and invasive disease, while loss of CPS biosynthesis and mucoidy can improve persistence during UTI. Combined, these data suggest that mucoidy may provide niche-specific advantages or disadvantages.

We hypothesized that environmental signals may control *K. pneumoniae* adaptation to the urinary tract. We found that hypervirulent, *rmp*-encoding strains and non-mucoid, *rmp-*negative strains have significantly reduced mucoidy when cultured in urine. We performed a transposon screen to identify genetic factors through which urine suppresses mucoidy in KPPR1. We collected 45 distinct transposon insertion isolates with insertions in 14 genes. Perplexingly, isolates with the same transposon insertion site exhibited different mucoidy phenotypes. Whole genome sequencing revealed that six distinct secondary point mutations in Wzc were present in every isolate with increased mucoidy and this corresponded with increased cell-free extracellular polysaccharide production (EPS). Over-expressing each Wzc *in trans* in wildtype KPPR1 increased mucoidy for five of the six point mutations, suggesting that five of the mutations exert a dominant phenotype over wildtype Wzc. Contrary to previous reports, none of the mutant Wzc over-expression strains increased CPS or EPS production, suggesting that the increased mucoidy is not due to overproduction of CPS or dispersal of CPS into the extracellular environment.^24,25^ One periplasmic mutation increased Wzc autophosphorylation, while four of the point mutations predicted to be in the Wzc active site reduce Wzc autophosphorylation. These same four point mutations are sufficient to increase the mucoidy of an *rmp-*negative clinical UTI isolate. Altogether, these data implicate Wzc activity in controlling *K. pneumoniae* mucoidy independent of *rmp*-regulation and increased CPS biosynthesis. Wzc activity may represent the lynchpin between mucoidy and CPS biosynthesis.

## RESULTS

### Urine suppresses mucoidy without reducing CPS biosynthesis

To examine the role of the urinary tract environment in *K. pneumoniae* mucoidy and CPS production, four strains were cultured in rich medium (LB) or sterile-filtered human urine. We selected two hypervirulent laboratory strains, KPPR1 (rifampin-resistant ATCC 43816 derivative) and NTUH-K2044, and two clinical UTI isolates, 616 and 1346.^26-28^ 616 and 1346 were originally identified as *K. pneumoniae* by MALDI-TOF, but we analyzed their genomes using *Pathogenwatch* and determined that, although both belong to the *K. pneumoniae* complex, 616 is *Klebsiella variicola* and 1346 is *Klebsiella quasipneumoniae*. This analysis determined that 616 encodes KL114 with an unknown K type and is predicted to produce O3/O3a LPS, while 1346 is predicted to produce K1 capsular polysaccharide (CPS) and O3/O3a LPS.^29-31^ Neither strain carries the *rmpADC*, genes known to regulate CPS biosynthesis and mucoidy.^16,17^ As expected, the UTI isolates were significantly less mucoid than the hypervirulent strains (**Fig. 1A**). Surprisingly, all four strains were significantly less mucoid in urine than in LB medium, suggesting that either LB medium activates or urine suppresses mucoidy (**Fig. 1A**). The bacterial cells were washed with PBS and the uronic acid content of cell-associated CPS was quantified. Culturing the strains in urine did not significantly impact CPS production as measured by uronic acid content, except it appears to stimulate CPS production in strain 616 (**Fig. 1B**). These results demonstrate that environmental conditions may regulate mucoidy independently of CPS and that mucoidy may be suppressed during UTI.

**Figure 1.**
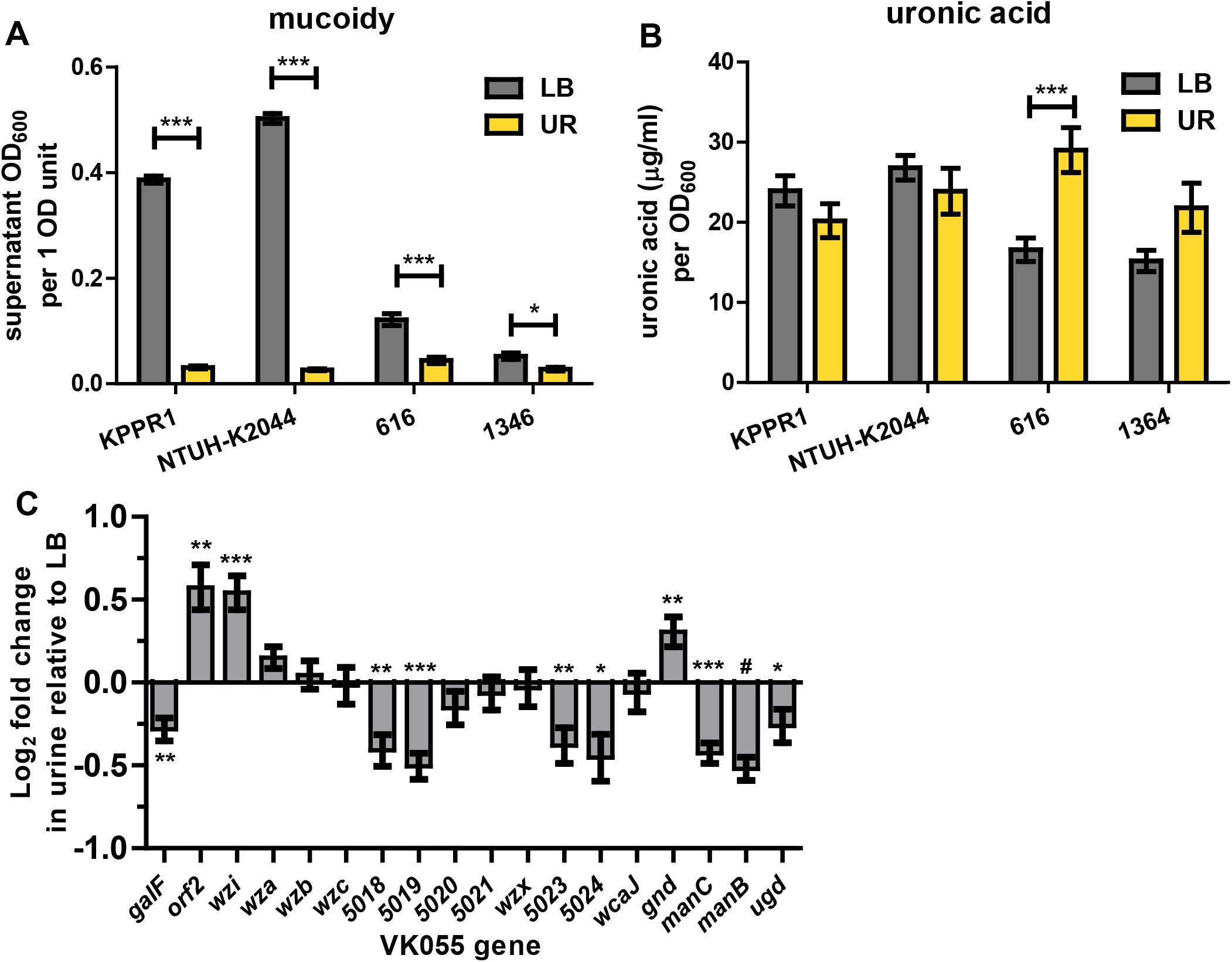
Urine suppresses mucoidy but not CPS biosynthesis. *K. pneumoniae* complex strains KPPR1, NTUH-K2044, 616, and 1364 were cultured in LB medium or sterile-filtered human urine (UR). (**A**) Mucoidy was determined by quantifying the supernatant OD600 after sedimenting 1 OD600 unit of culture at 1,000 x *g* for 5 min. (**B**) The uronic acid content of crude CPS extracts was determined after washing 1 OD600 unit of bacterial cells in PBS. (**C**) In addition, the relative abundance of each gene in the CPS biosynthesis operon in UR relative to LB medium was determined by qRT-PCR and normalized to *gap2* transcript abundance. Statistical significance in **A** and **B** was determined using two-way ANOVA with a Bonferroni post-test to compare specific groups. In **C**, a student’s *t* test was used to determine if each value was significantly different from 1.0. * *p* < 0.05; ** *p* < 0.01; *** *p* < 0.001; # *p* < 0.0001. All experiments were performed ≥3 independent times, in triplicate.

To examine CPS biosynthesis by another measure, we quantified the transcript abundance of each gene in the KPPR1 CPS biosynthetic gene cluster in urine relative to LB medium. CPS biosynthetic genes were not broadly suppressed in urine compared to LB medium (**Fig. 1C**). Rather, eight genes (*galF, VK055_5018, VK055_5019, VK055_5023, VK055_5024, manC, manB, ugd*) were significantly down-regulated, although the fold-change was minor (range = 0.704 - 0.850). Three genes (*orf2, wzi, gnd*) were significantly up-regulated, again with minor fold-change increase (range = 1.256 – 1.545). These fold changes do not impact the amount of CPS produced, as measured by uronic acid content in crude CPS extracts (**Fig. 1B**). Although these transcriptional changes could be responsible for reducing the mucoidy of bacterial culture, the small fold-change suggested that other factors may drive the suppression of mucoidy by urine.

### Medium alkalinity increases mucoidy and cell-free extracellular polysaccharides production

Since urine often has a pH lower than LB medium, we hypothesized that the pH of the medium could affect mucoidy or extracellular polysaccharide (EPS) production and localization. To discriminate between cell-associated EPS (CPS) versus cell-free EPS, we quantified the total uronic acid content of EPS extracted from both bacteria and culture medium (total EPS) versus the spent culture supernatant (cell-free EPS). CPS was inferred by subtracting cell-free EPS from total EPS. This approach circumvented technical issues caused by poor sedimentation, which confounds direct measurement of cell-associated CPS as it is challenging to cleanly separate cells from supernatant. We cultured KPPR1 in LB medium adjusted to pH 5, 6, 7, 8, and 9 then quantified mucoidy by sedimentation, and EPS localization by uronic acid content (**Fig. 2**). We observed that pH 9 increases mucoidy and cell-free EPS production, while pH 5 reduces mucoidy and cell-free EPS production. However, these changes in mucoidy are not as dramatic as what was observed in urine (**Fig. 1A-B**) and did not occur in the pH range between urine (pH 6.5) and LB medium (pH 7). These data indicate that basic pH increases both mucoidy and cell-free EPS, but that is not likely the factor by which urine suppresses mucoidy. It is possible that pH impacts mucoidy by altering cell-free EPS.

**Figure 2.**
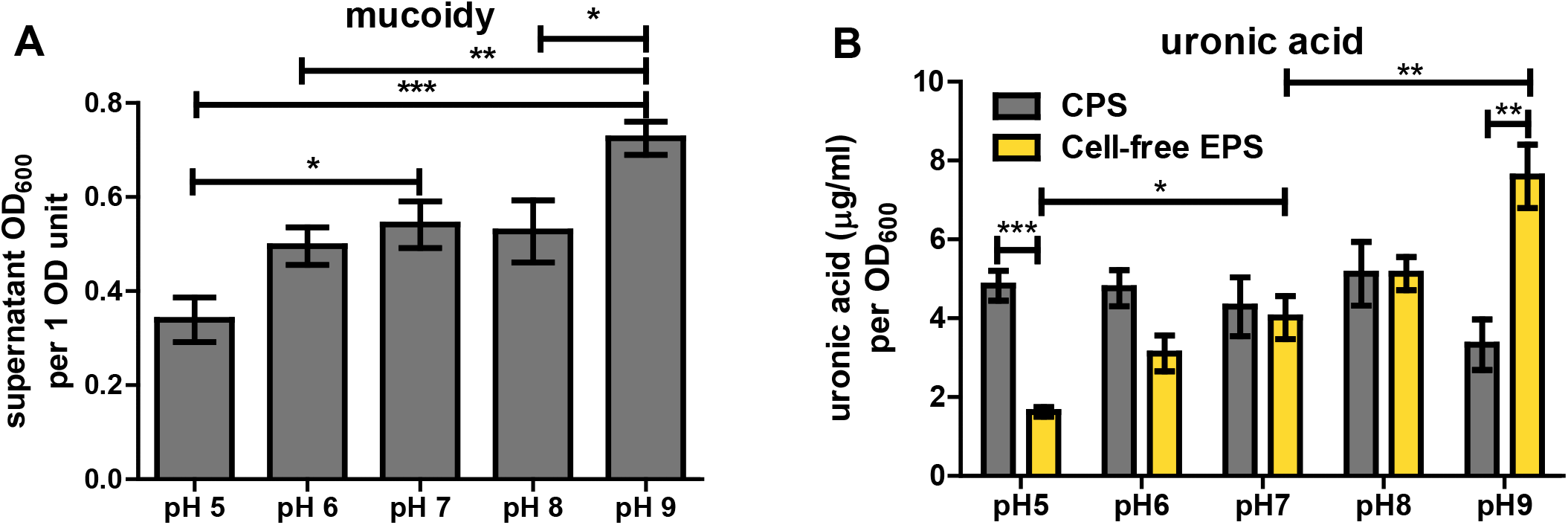
Changes in pH drives mucoidy and cell-free EPS production. *K. pneumoniae* KPPR1 was cultured in pH-adjusted LB medium. (**A**) Mucoidy was determined by quantifying the supernatant OD600 after sedimenting 1 OD600 unit of culture at 1,000 *x g* for 5 min. (**B**) EPS was extracted from either total culture or spent medium and the uronic acid content was determined and normalized to the OD600 of the overnight culture. Statistical significance was determined using two-way ANOVA with a Bonferroni post-test to compare specific groups. * *p* < 0.05; ** *p* < 0.01; *** *p* < 0.001; # *p* < 0.0001. Experiments were performed ≥2 independent times, in triplicate.

### A transposon screen for genes required urine to reduce mucoidy

To identify bacterial factors that can circumvent urine-suppressed mucoidy, we screened an arrayed KPPR1 transposon library for mutants with increased mucoidy in urine. The transposon library contains 3,733 mutants, which disrupts 72% of all open reading frames.^18^ The library was cultured in sterile, pooled human urine and the mucoidy of each mutant was assessed by sedimentation. With a hit rate of 2.4%, the 89 mutants initially identified to have increased mucoidy were evaluated for mucoidy by string test on urine agar and sedimentation efficiency in triplicate. 18 of 89 primary hits appeared mucoid either on urine agar or in the secondary sedimentation assay performed in triplicate. We noted that some hits had mixed colony morphology. Therefore, the string test on urine agar was repeated for each isolate. Ultimately, 14 transposon mutants had at least one isolate that passed all screening and confirmation (**Table 1**).

**Table 1.** Tn:mucoid^WT^ and Tn:mucoid^+^ isolate phenotypes and genotypes.

| Table 1. Tn:mucoi<sup>WT</sup> and Tn:mucoi<sup>+</sup> isolate phenotypes and genotypes. |  |  |  |  |  |  |  |  |
| --- | --- | --- | --- | --- | --- | --- | --- | --- |
| Clone ID | Gene | Primary screen: Sedimentation | Secondary screen: String test | Secondary screen: Sedimentation | Individual isolate: String test | WGS Non-synonymous Variants | Wzc Sanger Sequencing | Wzc residue |
|  | all transposon average | 0.0520 (+/- 0.0047) | 0 | .05275 (+/- .0046) | 0 |  |  |  |
|  | VK055_4579_pheA | 0.105 | + | 0.068 |  |  |  |  |
| 1 B1 | VK055_4579_pheA |  |  |  | + | VK055_5017_wzc C1939T | VK055_5017_wzc C1939T | WzcP646S |
| 2 B1 | VK055_4579_pheA |  |  |  | 0 | VK055_2438 C397T<br>VK055_4590 A34C |  |  |
| 3 B1 | VK055_4579_pheA |  |  |  | 0 | none reported |  |  |
|  | VK055_3922_gmhA2 |  | + | 0.069667 |  |  |  |  |
| 1 F2 | VK055_3922_gmhA2 |  |  |  | + |  |  |  |
| 1 F2g | VK055_3922_gmhA2 |  |  |  | + |  | VK055_5017_wzc C1183A | WzcQ395K |
| 2 F2 | VK055_3922_gmhA2 |  |  |  | + |  |  |  |
| 3 F2 | VK055_3922_gmhA2 |  |  |  | + |  | VK055_5017_wzc C1183A | WzcQ395K |
|  | Tn+, insertion unknown | 0.192 | + | 0.052333 |  |  |  |  |
| 4 B4 | Tn+, insertion unknown |  |  |  | + | VK055_1978_bepE T659G | VK055_5017_wzc G1709T | WzcG569V |
| 5 B4 | Tn+, insertion unknown |  |  |  | + | VK055_5017_wzc G1709T<br>VK055_0186_vgrG2 G2248A<br>VK055_1734 A212C | WT and<br>VK055_5017_wzc G1709T | WzcG569V |
| 6 B4 | Tn+, insertion unknown |  |  |  | 0 | none reported |  |  |
|  | VK055_1828 | 0.06 | + | 0.051667 |  |  |  |  |
| 7 C8 | VK055_1828 |  |  |  | 0 | VK055_5031 T406A |  |  |
| 8 C8 | VK055_1828 |  |  |  | 0 | VK055_4839 1382ΔG<br>intergenic 3A938201G |  |  |
| 9 C8 | VK055_1828 |  |  |  | + | VK055_5017_wzc G1709T | VK055_5017_wzc G1709A | WzcG569D |
|  | VK055_2345 | 0.08 | + | 0.053333 |  |  |  |  |
| 7 C9 | VK055_2345 |  |  |  | + |  | VK055_5017_wzc C1183A | WzcQ395K |
| 8 C9 | VK055_2345 |  |  |  | + |  | VK055_5017_wzc C1183A | WzcQ395K |
| 9 C9 | VK055_2345 |  |  |  | + |  | VK055_5017_wzc C1183A | WzcQ395K |
|  | VK055_4105 | 0.061 | + | 0.052333 |  |  |  |  |
| 7 C10 | VK055_4105 |  |  |  | + |  | VK055_5017_wzc G1708T | WzcG569C |
| 8 C10 | VK055_4105 |  |  |  | + |  | VK055_5017_wzc G1708T | WzcG569C |
| 9 C10 | VK055_4105 |  |  |  | + |  | VK055_5017_wzc G1708T | WzcG569C |
|  | VK055_0204 | 0.054 | + | 0.067333 |  |  |  |  |
| 7 D8 | VK055_0204 |  |  |  | + | VK055_5017_wzc C1939T<br>VK055_4651_pbpC 2033T>G<br>VK055_4651_pbpC 2045C>G | VK055_5017_wzc C1939T | WzcP646S |
| 8 D8 | VK055_0204 |  |  |  | 0 | VK055_4839 1382ΔG<br>intergenic C2225958T<br>intergenic C2225976A |  |  |
| 9 D8 | VK055_0204 |  |  |  | + | VK055_5017_wzc C1939T | VK055_5017_wzc C1939T | WzcP646S |

| Table 1 cont'd. Tn:mucoi<sup>WT</sup> and Tn:mucoi<sup>+</sup> isolate phenotypes and genotypes. |  |  |  |  |  |  |  |  |
| --- | --- | --- | --- | --- | --- | --- | --- | --- |
| Clone ID | Gene | Primary screen: Sedimentation | Secondary screen: String test | Secondary screen: Sedimentation | Individual isolate: String test | WGS Non-synonymous Variants | Wzc Sanger Sequencing | Wzc residue |
|  | all transposon average | 0.0520 (+/- 0.0047) | 0 | .05275 (+/- .0046) | 0 |  |  |  |
|  | VK055_1812_dtpD | 0.077 | + | 0.051333 |  |  |  |  |
| 7 E2 | VK055_1812_dtpD |  |  |  | + |  | VK055_5017_wzc WT+G1709A | Wzc<sub>WT+G569D</sub> |
| 8 E2 | VK055_1812_dtpD |  |  |  | + |  | VK055_5017_wzc WT+G1709A | Wzc<sub>WT+G569D</sub> |
| 9 E2 | VK055_1812_dtpD |  |  |  | + |  | VK055_5017_wzc G1708T | Wzc<sub>G569C</sub> |
|  | VK055_4261 | 0.075 | + | 0.050667 |  |  |  |  |
| 7 E3 | VK055_4261 |  |  |  | + |  | VK055_5017_wzc G1708A | Wzc<sub>G569D</sub> |
| 8 E3 | VK055_4261 |  |  |  | + |  | VK055_5017_wzc G1708A | Wzc<sub>G569D</sub> |
| 9 E3 | VK055_4261 |  |  |  | + |  | VK055_5017_wzc G1708A | Wzc<sub>G569D</sub> |
|  | VK055_0874 | 0.064 | + | 0.052 |  |  |  |  |
| 7 E5 | VK055_0874 |  |  |  | + |  | VK055_5017_wzc G1708A | Wzc<sub>G569D</sub> |
| 8 E5 | VK055_0874 |  |  |  | + |  | VK055_5017_wzc G1708A | Wzc<sub>G569D</sub> |
| 9 E5 | VK055_0874 |  |  |  | + |  | VK055_5017_wzc G1708A | Wzc<sub>G569D</sub> |
|  | VK055_4849_nuoN |  | + | 0.075 |  |  |  |  |
| 7 F7 | VK055_4849_nuoN |  |  |  | 0 | VK055_4892_mqo2 T1289G<br>VK055_0435 C587T |  |  |
| 8 F7 | VK055_4849_nuoN |  |  |  | + | VK055_5017_wzc G1709T | VK055_5017_wzc G1709T | Wzc<sub>G569V</sub> |
| 9 F7 | VK055_4849_nuoN |  |  |  | + | none reported | VK055_5017_wzc G1709T | Wzc<sub>G569V</sub> |
|  | VK055_4923_pfkB | 0.059 | + | 0.050667 |  |  |  |  |
| 7 F9 | VK055_4923_pfkB |  |  |  | 0 | none reported | VK055_5017_wzc T798C | synonymous |
| 7 F9g | VK055_4923_pfkB |  |  |  | + | VK055_5017 1576_insGCA_1577 | VK055_5017 1576_insGCA_1577 | VK055_5017 526_insA_527 |
| 8 F9 | VK055_4923_pfkB |  |  |  | 0 | VK055_0184 A1007T |  |  |
| 9 F9 | VK055_4923_pfkB |  |  |  | 0 | none reported |  |  |
|  | VK055_2280_proB | 0.065 | + | 0.066333 |  |  |  |  |
| 7 H8 | VK055_2280_proB |  |  |  | 0 | VK055_3855 G211T<br>intergenic C879584A |  |  |
| 8 H8 | VK055_2280_proB |  |  |  | 0 | none reported |  |  |
| 9 H8 | VK055_2280_proB |  |  |  | + | VK055_5017_wzc G1709T<br>VK055_1078 G586T | VK055_5017_wzc WT+G1709A+G17 | Wzc<sub>WT+G569D+G569V</sub> |
|  | VK055_1417 | 0.069 | + | 0.055667 |  |  |  |  |
| 10 D5 | VK055_1417 |  |  |  | 0 |  |  |  |
| 10 D5g | VK055_1417 |  |  |  | + |  | VK055_5017_wzc C1183A | Wzc<sub>Q395K</sub> |
| 11 D5 | VK055_1417 |  |  |  | + |  | VK055_5017_wzc C1183A | Wzc<sub>Q395K</sub> |
| 11 F10 | VK055_1417 |  |  |  | + |  | VK055_5017_wzc C1183A | Wzc<sub>Q395K</sub> |

Elevated CPS production is associated with increased mucoidy, although there is strong evidence that other factors contribute to mucoidy.^17,18^ Each of the 14 transposon mutants had 3 or 4 isolates that exhibited either wildtype (Tn:mucoid^WT^) or increased mucoidy (Tn:mucoid^+^) (N = 45 isolates total) (**Table 1**). Each isolate was cultured in triplicate and assayed for total extracellular uronic acid content. On average, the Tn:mucoid^+^ isolates produce 1.15-fold more extracellular uronic acid compared to Tn:mucoid^WT^ isolates (**Fig. 3**). This difference is modest, but significant. Notably, the two Tn:mucoid^WT^ isolates that produced the most uronic acid have transposon insertions in *gmhA2*. All four *gmhA2* transposon mutants exhibit elevated uronic acid production (mean = 11.89 μg/mL uronic acid per OD_600_), regardless of mucoidy state (**Fig. 3**, open markers). It is also notable that most Tn:mucoid^+^ strains produce extracellular uronic acid quantities comparable to Tn:mucoid^WT^ strains (**Fig. 3**, blue band), adding further evidence that increased secretion of extracellular glucuronidated polysaccharides is not the only factor driving *K. pneumoniae* mucoidy.

**Figure 3.**
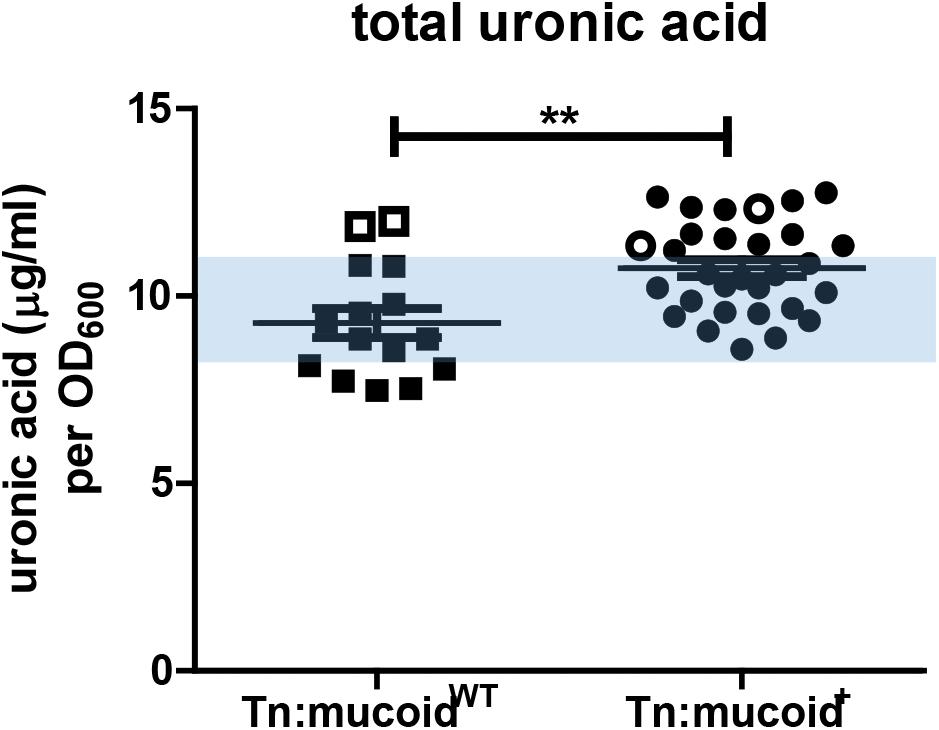
Transposon isolates with elevated mucoidy produce more extracellular uronic acid. Each *K. pneumoniae* transposon isolate was cultured in triplicate in LB medium, where N = 45 isolates representing 14 transposon strains. Extracellular polysaccharides were extracted from total culture and the uronic acid content was determined and normalized to the OD600 of the overnight culture. Each marker represents an individual isolate. Open markers identify *gmhA2* transposon insertion mutant isolates. A student’s *t* test was used to determine if the mean extracellular uronic acid produced in strains that exhibit elevated mucoidy (Tn:mucoid^+^) was significantly different compared to transposon strains that exhibited wildtype mucoidy (Tn:mucoid^WT^), where ** *p* < 0.01.

### Transposon insertion sites are validated by PCR

The isolate-to-isolate variation in increased mucoidy was puzzling. The phenotype of each isolate was stable after passaging. The transposon insertion site of each isolate was verified by PCR with one primer that anneals to the transposon and one primer that anneals to the expected genomic region. The B4 transposon strain was selected from a well that did not have the insertion site identified according to our previous cartesian pooling and coordinate sequencing protocol.^18,32^ In case it arose as cross-contamination with another hit, we tested the colonies for amplification with all other sets of PCR primers, but did not get any amplification. We did get amplification with primers that only annealed within the transposon, indicating the presence of a transposon at an unidentified site.

### Tn:mucoid^+^ strains contain secondary mutations in wzc

Since most hits had the expected transposon insertion site but exhibited varying mucoidy, even within a single transposon mutant, we hypothesized that a secondary point mutation was driving the increased mucoidy. We submitted purified genomic DNA from 22 isolates (N = 10 Tn:mucoid^+^ and N = 12 Tn:mucoid^WT^), plus our wildtype KPPR1, for whole genome sequencing (WGS) (**Table 1**).

Sequence variants in each strain were identified using the Variation Analysis pipeline on PATRIC using *K. pneumoniae* subsp. *pneumoniae* strain VK055 (RefSeq: <u>NZ_CP009208.1</u>) as the reference strain.^33,34^ Comparisons between our laboratory KPPR1 and the published genome identified 1 non-synonymous (insG) mutation at position 1610890 in *VK055_1597* and a synonymous (C>T) mutation at position 2880686 in *VK055_2811*. These variants were disregarded in all transposon isolates. 31 other variants were identified (N = 2 deletions, N = 1 insertion, N = 20 nonsynonymous, N = 4 intergenic, and N = 4 synonymous). The one insertion and seven nonsynonymous mutations all mapped to *VK055_5017_etk* (*wzc*) and were associated with eight of the ten Tn:mucoid^+^ isolates. No other genetic variations were associated with more than one Tn:mucoid^+^ isolate. Six unique variations associated with Tn:mucoid^+^ isolates included, *VK055_0186_vgrG2*_*2248G>A*_, *VK055_1078*_*586G>T*,_ *VK055_1734*_*212A>C*_, *VK055_1978_bepE*_*659T>G*_, *VK055_4651_pbpC*_*2033T>G*_, *and VK055_4651_pbpC*_*2045C>G*_ (**Table 1**). The remaining 13 variants were associated with Tn:mucoid^WT^ isolates (N = 7 of 12) and none had point mutations in *wzc*. 90% of Tn:mucoid^+^ had at least one non-synonymous variant, while 58% of Tn:mucoid^WT^ had at least one non-synonymous variant (**Fig. S1**), suggesting that the selection of Tn:mucoid^+^ strains enriched the number of secondary point mutations.

Since only 22 of the 45 transposon isolates were submitted for whole genome sequencing, the *wzc* gene of all 45 isolates was PCR amplified and the open reading frame interrogated for point mutations using Sanger sequencing. Sequences were aligned in DNASTAR LaserGene MegAlign using Clustal W. All strains had zero gaps and the chromatogram trace was used to resolve questionable base calls or the purity of the culture. All 30 Tn:mucoid^+^ had non-synonymous mutations in *wzc*, including the two isolates that did not have *wzc* mutations identified by WGS. None of the 16 Tn:mucoid^WT^ isolates had *wzc* mutations. In four instances, Tn:mucoid^+^ appeared to be a mixture of wildtype and a residue change at Wzc_G569_ based on chromatogram (5 B4, 7 E2, 8 E2, and 9 H8) (**Table 1**), indicating that the phenotype is dominant if the colonies are not fully purified. Twice WGS called *wzc*_G1709A_ and Sanger sequencing called *wzc*_G1709T_ (9 C8 and 9 H8) (**Table 1**), which could be due to heterogeneity at this base pair.

Six unique Wzc mutations were distributed across the 30 Tn:mucoid^+^ strains. The Wzc mutations resulted in the following amino acid changes, Wzc_Q395K_ (N = 8), Wzc_526insA_ (N = 1), Wzc_G569C_ (N = 6), Wzc_G569V_ (N = 4), Wzc_G569D_ (N = 8), and Wzc_P646S_ (N = 3). Wzc is a conserved bacterial tyrosine kinase involved in CPS biosynthesis.^19^ It forms a periplasmic octamer that interfaces with the Wza octamer in the outer membrane and the Wzb phosphatase in the periplasm.^20,21,35^ The current model is that Wzc autophosphorylation dissociates the complex, with subsequent Wzb-mediated dephosphorylation resulting in complex re-assembly.^21^ It is thought that these protein dynamics drive Wza extrusion of nascent extracellular polysaccharides, as loss of either Wzc phosphorylation or dephosphorylation disrupts CPS formation.^20^ Wzc is a two-membrane pass monomer with residues 1-30 and 448-722 located in the cytoplasm and residues 51-427 located in the periplasm. Only Wzc_Q395K_ is located in the periplasmic region in motif 3 (**Fig. 4**). Deletion of motif 3 in structural studies of *E. coli* Wzc resulted in loss of CPS production, with neither loss of octamerization nor loss of autokinase activity.^21^ Therefore, it is thought that motif 3 interfaces with Wza. The remaining point mutations are located in the C-terminal cytoplasmic region responsible for autokinase activity. Three Wzc_G569_ mutations were identified (G>C, G>V, and G>D) (**Fig. 4**). Wzc_G569_ is adjacent to Wzc_Y570_, which is the active reside that auto-phosphorylates the poly-tyrosine C-terminus. Wzc_P646S_ is located in the Walker B motif involved in ATP binding in the active site (**Fig. 4**).^36^ The alanine insertion after residue 526 is 8 residues up-stream of a Walker A motif (**Fig. 4**), which also contributes to active site ATP binding. Nonetheless, residue 526insA is not located in the active site and it is unclear if it is located in another domain critical for kinase activity or oligomerization.

**Figure 4.**
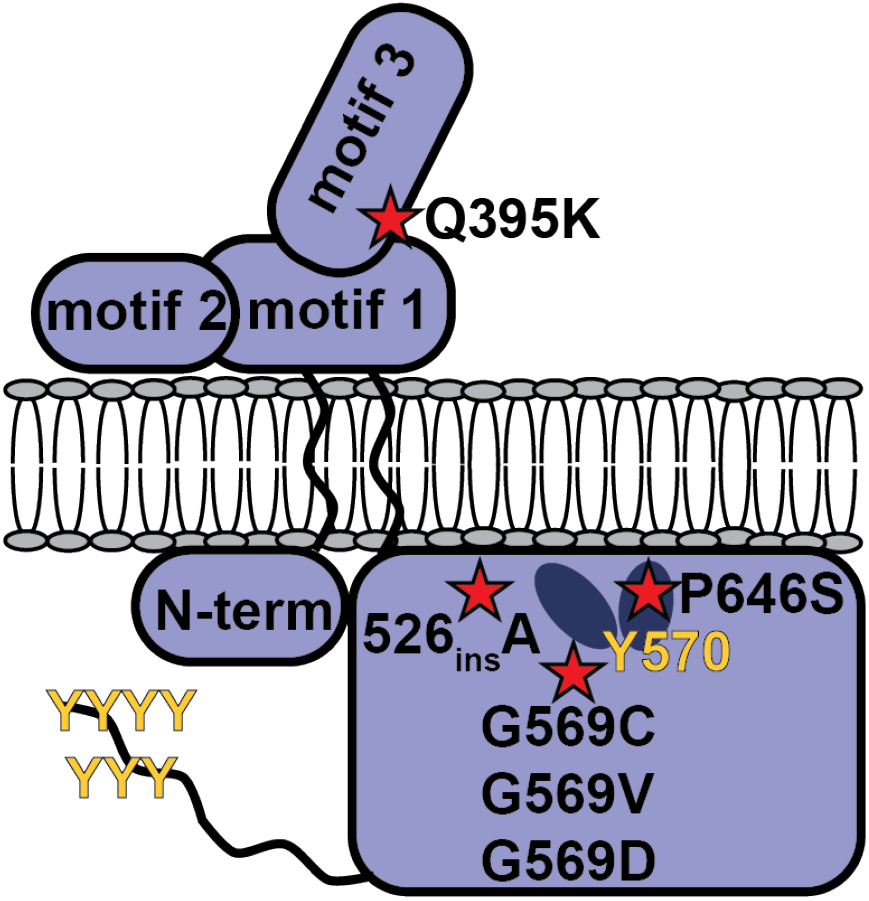
Location of Wzc point mutations in Tn:mucoid^+^ strains. Six unique Wzc point mutations were identified in all 30 Tn:mucoid^+^ strains. The Wzc mutations are marked with a red star, where the dark ovals represent Walker A and B motifs and the yellow Ys represent tyrosine residues involved in autophosphorylation. WzcQ395K is localized to the periplasmic motif 3, predicted to interact with the Wza outer membrane protein. WzcG569 is adjacent to the active site tyrosine. WzcP646S is located in the active site Walker B motif. Wzc526insA is located 8 residues up-stream of the active site Walker A motif.

### Tn:mucoid^+^ strains impact mucoidy and increase cell-free EPS levels

We selected at least one Tn:mucoid^+^ isolate representative of each of the six Wzc point mutations. We prioritized mutants based on if they had a partner Tn:mucoid^WT^ strain and if we had high confidence about their Wzc genotype. In all instances, the six Wzc mutations increased cell-free EPS production relative to wildtype KPPR1 and/or their Tn:mucoid^WT^ strain (**Fig. 5B**). Similarly, all Tn:mucoid^+^ appeared more mucoid based on sedimentation assay, except for Tn:mucoid^+^ strains encoding Wzc_Q395K_ (**Fig. 5A**). Since these strains exhibited a different phenotype from the other Wzc mutant strains, we assayed two transposon families that encoded Wzc_Q395K_, *gmhA2* (1 F2, 1F2g, 2 F2, and 3 F2) and *VK055_2345* (7 C9). All three isolates encoding Wzc_Q395K_ exhibited reduced mucoidy, based on sedimentation, despite overt mucoidy of the cultures (**Fig. 5**). Furthermore, the *gmhA2* Tn:mucoid^WT^ isolates alone have increased EPS production, which tends to make the strains appear more mucoid, although only 1 F2 has significantly increased mucoidy based on sedimentation. The *gmhA2-*Wzc_Q395K_ (1 F2g and 3 F2) isolates appear to have completely lost the ability to anchor CPS to the cell surface, resulting in all EPS being cell-free. It is possible the lack of any cell-associated CPS is why these strains appear visibly mucoid, but sediment well. Previous work has shown that some CPS is required for mucoidy to be apparent in the sedimentation assay.^17,18^

**Figure 5.**
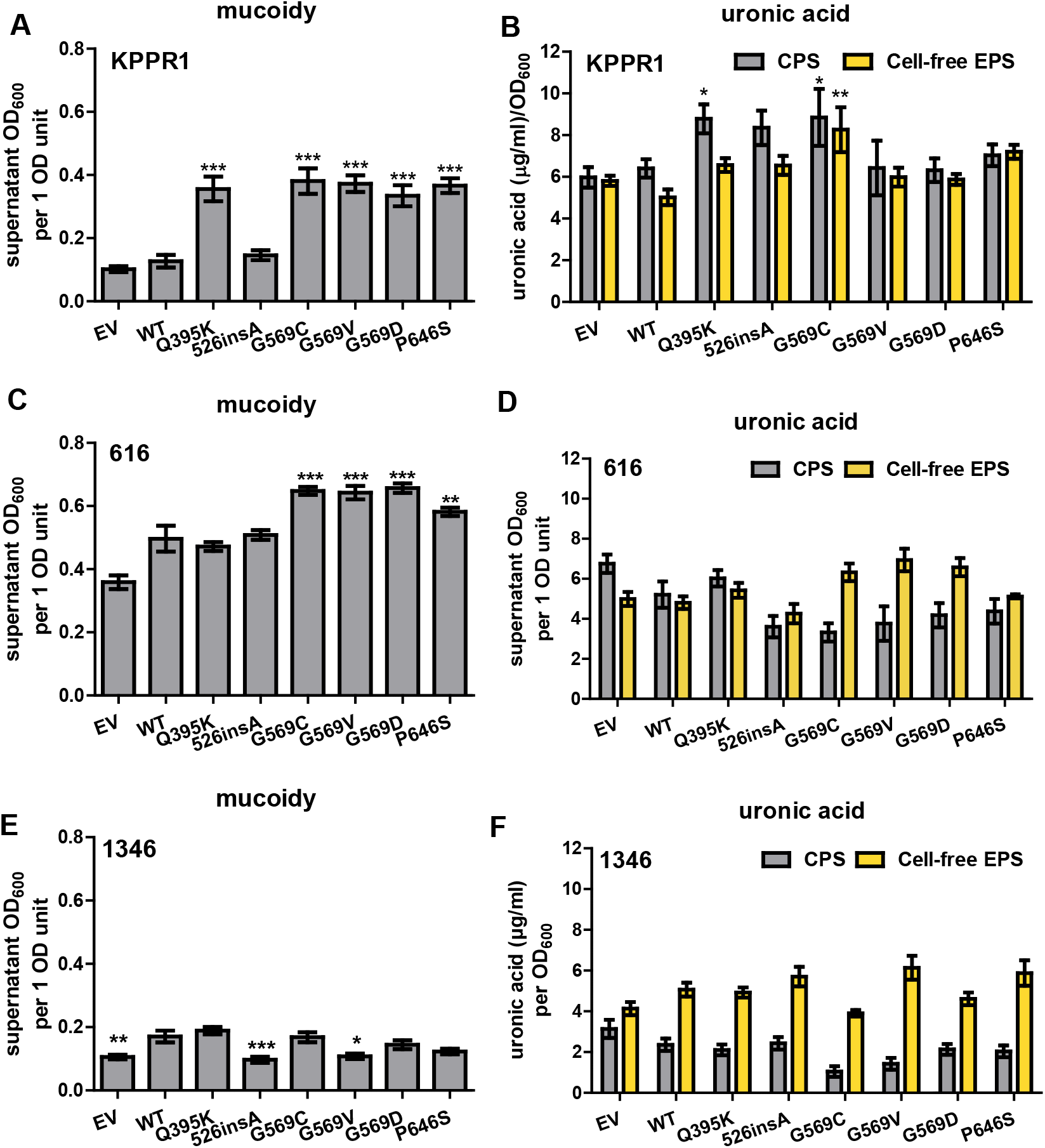
Tn:mucoid^+^ strains impact mucoidy and EPS production. *K. pneumoniae* Tn:mucoid^+^ or Tn:mucoid^WT^ strains were cultured in LB medium. The isolate identifier is listed on the X-axis along with the Wzc mutation listed in parentheses (WT = wildtype, QK = Q395K, insA = 526insA, GC = G569C, GV = G569V, GD = G569D, PS = P646S). Transposon insertion sites can be found in **Table 1**. (A) Mucoidy was determined by quantifying the supernatant OD600 after sedimenting 1 OD600 unit of culture at 1,000 *x g* for 5 min. (B) The uronic acid content of the total culture or supernatant were quantified and normalized to the OD600 of the overnight culture. The cell-associated CPS was deduced by subtracting the cell-free uronic acid content (EPS) from the total culture uronic acid content. Statistical significance was determined using two-way ANOVA with a Bonferroni post-test to compare specific groups. * *p* < 0.05; ** *p* < 0.01; *** *p* < 0.001; # *p* < 0.0001. Experiments were performed ≥3 independent times, in triplicate.

### Wzc mutations are sufficient to increase mucoidy without impacting CPS production

Since it became clear that the transposon insertion sites may also impact mucoidy and CPS, we introduced the Wzc mutations into wildtype KPPR1 on a pBAD18 plasmid under an arabinose-inducible promoter. With the exception of Wzc_526insA_, over-expressing each mutation in KPPR1 significantly increased mucoidy compared to over-expressing wildtype Wzc (**Fig. 6A**). Furthermore, over-expressing wildtype Wzc had no effect on mucoidy or EPS production compared to empty vector. This indicates that it is not the copy number of Wzc, but the mutant Wzc proteins alone that exert an effect on mucoidy, even with an endogenous copy of wildtype Wzc. However, most point mutations did not increase CPS or cell-free EPS production, with the exception of Wzc_Q395K_ and Wzc_G569C_ modestly increasing CPS (1.4-fold) and Wzc_G569C_ increasing cell-free EPS (1.6-fold) (**Fig. 6B**). This indicates that although the Tn:mucoid^+^ strains increase mucoidy and cell-free EPS, it is not cell-free EPS driving the increased mucoidy in the Tn:mucoid^+^ strains.

**Figure 6.**
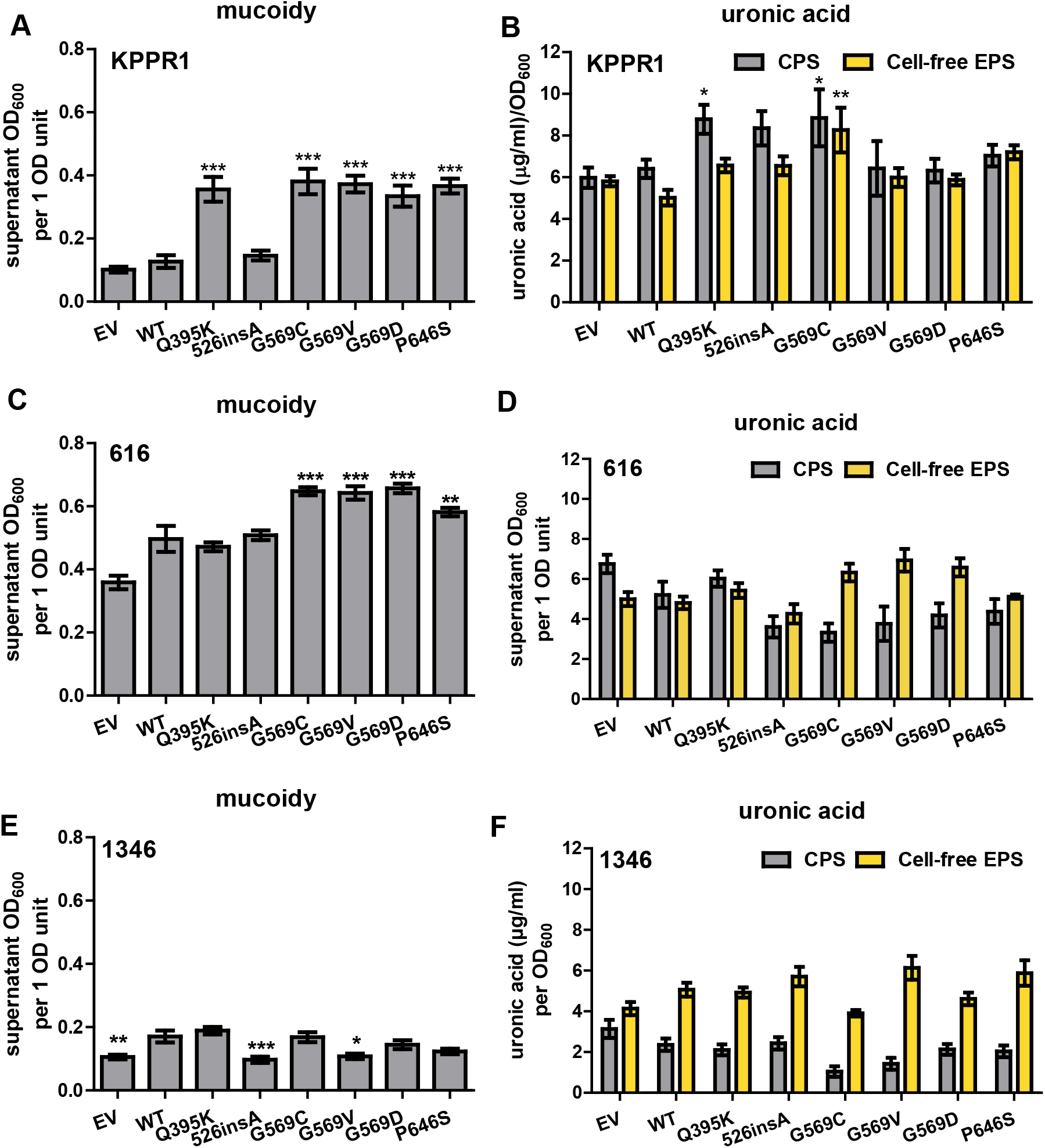
Over-expressing Wzc mutations increase mucoidy, but not EPS production. Wildtype Wzc and six mutants were expressed with the pBAD18 arabinose-inducible promoter in wildtype KPPR1 (**A-B**), 616 (**C-D**), or 1346 (**E-F**). Empty vector (EV) was used as a negative control. The *K. pneumoniae* strains were cultured in LB medium containing 50 mM L-arabinose and kanamycin. (A, C, E) Mucoidy was determined by quantifying the supernatant OD600 after sedimenting 1 OD600 unit of culture at 1,000 *x g* for 5 min. (B, D, F) Uronic acid content of the total culture or supernatant was quantified and normalized to the OD600 of the overnight culture. The cell-associated CPS was deduced by subtracting the cell-free uronic acid content (EPS) from the total culture uronic acid content. Statistical significance was determined using two-way ANOVA with a Bonferroni post-test to compare specific groups. * *p* < 0.05; ** *p* < 0.01; *** *p* < 0.001; # *p* < 0.0001. Experiments were performed ≥3 independent times, in triplicate.

We then examined if the defined Wzc mutations are sufficient to increase mucoidy in 616 and 1346. pBAD18 vectors carrying wildtype *wzc* or the six point mutants were transformed into 616 and 1346. Over-expressing Wzc_G569C_, Wzc_G569V_, Wzc_G569D_, and Wzc_P646S_ *in trans* is sufficient to increase mucoidy in 616, but not 1346 (**Fig. 6C** and **E**). CPS and cell-free EPS uronic acid content was not significantly different in the mutant Wzc strains compared to wildtype.

Altogether these data demonstrate that specific Wzc mutations are sufficient to increase mucoidy regardless of whether a strain encodes *rmpADC* (e.g. KPPR1) or not (e.g. 616). Furthermore, these data indicate that other cellular factors are required as the identified Wzc mutations are not sufficient to increase mucoidy in 1346. It is not likely the CPS type as 1346 encodes K1. The K1 CPS type is often associated with mucoid strains, but it is not sufficient for mucoidy as some K1 strains are non-mucoid.

### Wzc mutations alter cellular tyrosine phosphorylation

Since many of the identified Wzc mutations are predicted to be close the active site of this tyrosine kinase, we hypothesized that they may alter the cellular tyrosine phosphorylation profile. Whole cell lysates were separated by SDS-PAGE and their phospho-tyrosine (Y-P_i_) profile probed by western blot. As expected, we consistently observed Wzc Y-P_i_ at 80 kDa, possibly with a faint dimer at 160 kDa (**Fig. 7A, C**, and **E**). The 80 kDa band was confirmed to be Wzc Y-P_i_ by inducing expression of Wzc-His_6_ with arabinose (**Fig. S2A-B**). We also observed Y-P_i_ protein bands that migrated at approximately 15, 23, 27, and 42 kDa. These are not likely to be Wzc fragments as they do not react with His_6_-specific antibodies (**Fig. S2**). Furthermore, we confirmed that the Wzc-His_6_ variants exerted a similar effect on mucoidy as the untagged versions (**Fig. S2C** vs **Fig. 6A**).

**Figure 7.**
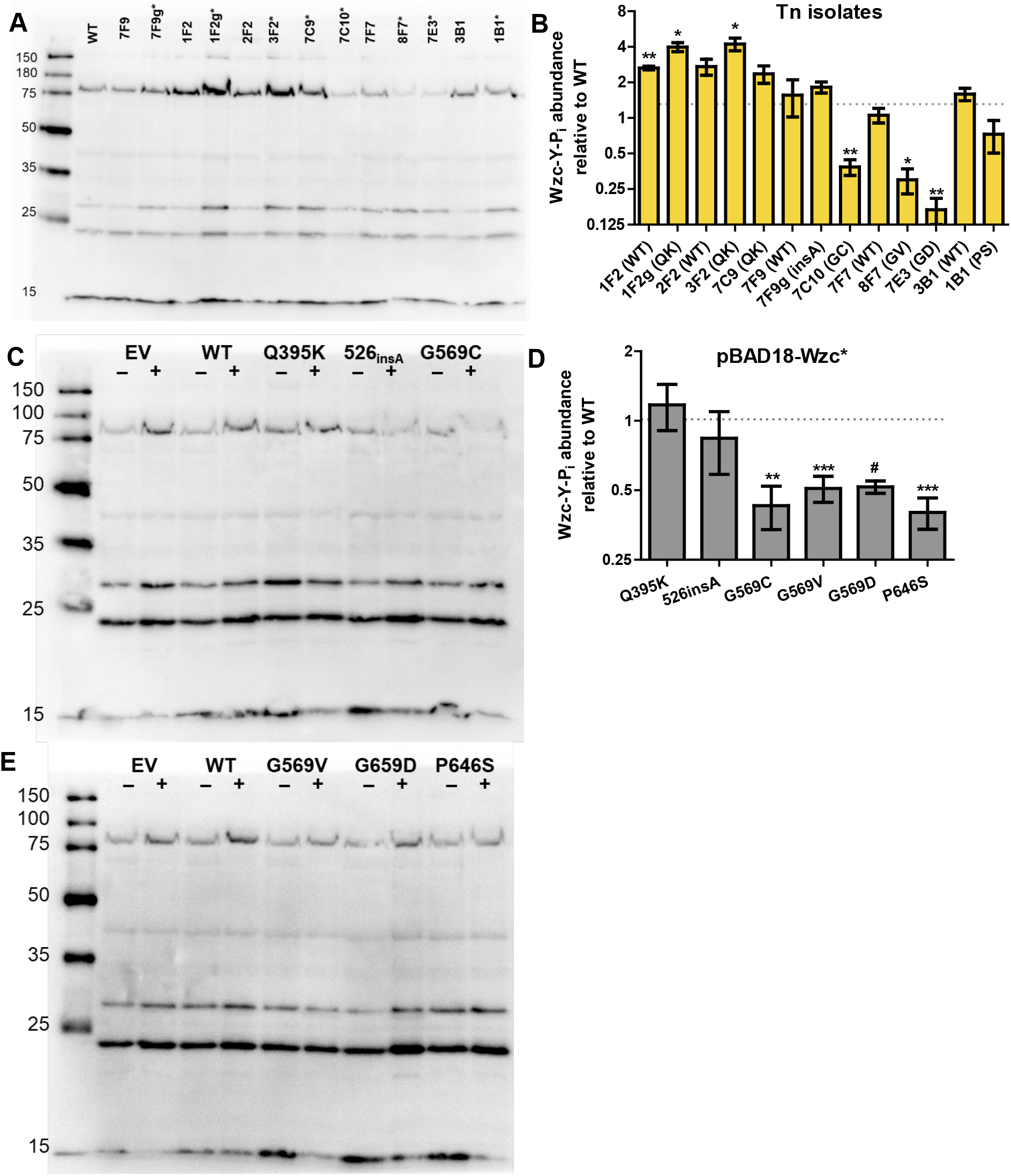
Anti-phosphotyrosine profiles of Wzc variants. Whole cell lysates prepared from *K. pneumoniae* Tn:mucoid^+^ or Tn:mucoid^WT^ strains cultured in LB medium were (**A**) resolved by SDS-PAGE, transferred to nitrocellulose, then probed with anti-phosphotyrosine antibody (PY20). (**B**) The Wzc-Y-Pi bands in **A** were then quantified in ImageJ and normalized to wildtype (WT) KPPR1 levels within each blot. (**C**-**D**) KPPR1 transformed with His6-tagged wildtype (WT) Wzc, six Wzc variants (Q395K, 526insA, G569C, G569V, G569D, P646S) or empty vector pBAD18 (EV) was cultured in LB medium containing kanamycin with (+) and without (-) 50 mM L-arabinose. Whole cell lysates were resolved by SDS-PAGE, transferred to nitrocellulose, then probed with antiphosphotyrosine antibody (PY20). (**E**) The Wzc-Y-Pi bands in **C-D** were quantified in ImageJ and normalized to KPPR1 over-expressing WT Wzc (lane 5) within each blot. A student’s *t* test was used to determine if each mean was significantly different from 1.0, (WT), where * *p* < 0.05; ** *p* < 0.01; *** *p* < 0.001; # *p* < 0.0001. In all instances lysates were prepared and analyzed ≥ 3 independent times. One representative image is shown.

We probed the panel of Tn:mucoid^+^ mutants and their partner Tn:mucoid^WT^ strains described in **Figure 5** with anti-phosphotyrosine antibody (PY20). We consistently observed increased Wzc Y-P_i_ phosphorylation in the transposon strains with *gmhA2* insertion sites (1 F2 and 2 F2), with the *gmhA2-*Wzc_Q395K_ (1 F2g and 3 F2) strains exhibiting further increases in Wzc Y-P_i_ abundance (**Fig. 7A-B**). 7 C9-Wzc_Q395K_ and 7 F9g-Wzc_526insA_ also tended to increase Wzc Y-P_i_ abundance compared to WT KPPR1 or 7 F9. The remaining Wzc point mutations (Wzc_G569C_, Wzc_G569V_, Wzc_G569D_, and Wzc_P646S_) appear to decrease Wzc Y-P_i_ abundance compared to their Tn:mucoid^WT^ counterpart strains or WT. With the exception of 7 F9 and 7 F9g-Wzc_526insA_, it appears that Wzc mutations that increase Wzc-Y-P_i_ compared to their Tn:mucoid^WT^ counterpart correlate with improved sedimentation, while Wzc mutations with relatively decreased Wzc-Y-P_i_ correlate with poor sedimentation (**Fig. 5A** and **7A-B**). Shifts in Wzc-Y-P_i_ abundance neither correlate with total of EPS produced nor the ratio of cell-free EPS to CPS, further emphasizing that other factors beyond CPS and EPS are required for mucoidy (**Fig. 5B** and **7A-B**). The 15, 23, and 27 kDa bands generally follow the same Y-P_i_ trends as Wzc, so it is possible that these bands represent other cellular factors that could control mucoidy. No changes were observed in the 42 kDa band, which suggests that this is non-specific.

We then probed wildtype KPPR1 carrying pBAD18 empty vector (EV), wildtype (WT) or each the six Wzc mutations with anti-phosphotyrosine antibody (PY20). Similar to the Tn:mucoid^+^ results, inducing the expression of Wzc_G569C_, Wzc_G569V_, Wzc_G569D_, and Wzc_P646S_ appear to decrease Wzc Y-P_i_ abundance compared to Wzc_WT_, while Wzc_Q395K_ and Wzc_526insA_ Y-P_i_ abundance remains similar to Wzc_WT_ (**Fig. 7C-E**). All pBAD18 vectors over-expressing Wzc mutations exhibited increased mucoidy compared to wildtype, except for Wzc_526insA_. (**Fig. 6A**). Therefore, it is likely that the Wzc_G569C_, Wzc_G569V_, Wzc_G569D_, and Wzc_P646S_ mutations affect mucoidy and Wzc-Y-P_i_ abundance by one mechanism, while the Wzc_Q395K_ and Wzc_526insA_ mutations affect mucoidy by other mechanisms. The 15, 23, and 27 kDa had variable signal intensity in several strains, though no discernable pattern with specific Wzc mutations was apparent (**Fig. 7A, C-D**). The identity of these other Y-P_i_ proteins is unknown. It was also apparent that culturing the bacteria with L-arabinose increased the cellular Y-P_i_ abundance even in KPPR1 carrying empty vector pBAD18 (**Fig. 7C-D**). We noted that treatment of KPPR1 with L-arabinose reduced mucoidy (**Fig. 5A** and **6A**), which may represent another exogenous signal that regulates mucoidy. Despite L-arabinose suppressing mucoidy, overexpressing Wzc_Q395K_, Wzc_G569C_, Wzc_G569V_, Wzc_G569D_, and Wzc_P646S_ using L-arabinose increased mucoidy, suggesting that these mutations function down-stream of L-arabinose signaling (**Fig. 6A**).

## DISCUSSION

Here, we have found that *K. pneumoniae* complex strains cultured in urine are significantly less mucoid than when cultured in LB medium, including strains that do not encode *rmpADC* (**Fig. 1A**). However, comparable levels of CPS production are sustained in both environments (**Fig. 1B**). This supports recent data demonstrating that mucoidy and CPS can be independently regulated, and extends these data to suggest that independent regulation can occur in response to discrete environmental signals.^16,17,37^ Although we have not yet identified those environmental signals present in LB medium versus urine, others have recently reported that L-fucose activates mucoidy independent of CPS and we have noted here that L-arabinose suppresses mucoidy independent of CPS.^38^ Therefore, it remains possible that either LB medium provides an environment supportive of mucoidy or urine contains signals that suppress mucoidy. We considered the possibility that the lower pH of urine (pH 6.5) versus LB medium (pH 7) could drive changes in mucoidy. We found that acidic pH reduces mucoidy and cell-free EPS while alkaline conditions boost mucoidy and cell-free EPS (**Fig. 2**). Although, pH does impact mucoidy, its effect is insignificant in the small pH difference between LB medium and urine.

We screened a transposon library to identify genetic factors that could restore mucoidy when bacteria are cultured in urine. We identified 45 distinct isolates representing 14 transposon insertion sites that increase mucoidy on urine (**Table 1**). Inconsistent mucoidy phenotypes within a single transposon insertion strain suggested that secondary mutations may be responsible for increasing mucoidy on urine. Whole genome sequencing with subsequent targeted Sanger sequencing determined that all isolates with increased mucoidy had six distinct secondary point mutations in the Wzc tyrosine kinase (**Fig. 4**). These Wzc mutations appear to fall into three functional classes that behave distinctly: (1) active site localized (Wzc_G569C_, Wzc_G569V_, Wzc_G569D_, Wzc_P646S_), (2) cytoplasmic (Wzc_526insA_), (3) periplasmic (Wzc_Q395K_). Only one transposon insertion mutant (*gmhA2*) had increased mucoidy in two isolates with wildtype *wzc* (**Fig. 5A**). The *gmhA2* transposon insertion strain also exhibited increased cell-free EPS (**Fig. 5B**). On average, *gmhA2* transposon insertion isolates had 2.7-fold greater Wzc-Y-P_i_ abundance compared to wildtype KPPR1 (**Fig. 7A-B**).

GmhA2 is predicted to associate with DnaA, initiating chromosome replication. It contains two sugar isomerase (SIS) domains, which bind phosphosugars. GmhA2 has homology to the GmhA phosphoheptose isomerase required for LPS biosynthesis, but has extended N- and C-terminal residues. We predict that GmhA2 regulates DNA replication in response to specific intracellular phosphosugars. BLASTP analysis of GmhA2 returns 674 homologs completely restricted to the Proteobacteria, with 672 homologs in Enterobacterales, 1 homolog in *Pseudomonas aeruginosa*, and 1 homolog in *Delftia acidovorans*. The GmhA binding partner, GmhB, has recently been shown to be critical for *K. pneumoniae* dissemination to the spleen via the bloodstream.^39^ No reports on the function of GmhA2 have been published and one can only speculate how this proteins regulates *K. pneumoniae* CPS production and mucoidy. It is certainly intriguing how a protein predicted to bind phosphosugars and regulate chromosomal replication alters Wzc-Y-P_i_, EPS anchoring to the cell surface, and mucoidy.

Two of the four *gmhA2* transposon mutants acquired a Wzc_Q395K_ secondary mutation (**Table 1**). *gmhA2* switched from producing more cell-free EPS to only producing an over-abundance of cellfree EPS when the secondary Wzc_Q395K_ mutation was acquired (**Fig. 5B**). Although the *gmhA2-* Wzc_Q395K_ appeared hypermucoid by string test and visual inspection, these strains surprisingly sedimented more efficiently than wildtype KPPR1 or *gmhA2* (**Fig. 5A**). In addition, acquisition of Wzc_Q395K_ in *gmhA2* further increased Wzc-Y-P_i_, 1.5-fold over *gmhA2-*Wzc_WT_ and 4.1-fold versus wildtype KPPR1. The Wzc_Q395K_ mutation was also isolated in the background of *VK055_2345* and *VK055_1417*. We further characterized the *VK055_2345-*Wzc_Q395K_ and found that it too exhibited sedimented more efficiently despite overt colony tackiness, increased cell-free EPS production, and 2.4-fold increase in Wzc-Y-P_i_ abundance compared to wildtype KPPR1 (**Table 2**). These phenotypes were not as dramatic as the *gmhA2-*Wzc_Q395K_ isolates, implicating a functional interaction between GmhA2 and Wzc in controlling mucoidy, EPS production, and tyrosine kinase activity. Overexpressing Wzc_Q395K_ *in trans* in wildtype KPPR1 increases mucoidy both visually and by sedimentation assay; increases CPS production, but not cell-free EPS; and does not change Wzc-Y-P_i_ abundance compared to wildtype KPPR1 (**Table 2**).

**Table 2.** Summary of Wzc Mutant Phenotypes.

| Table 2. Summary of Wzc Mutant Phenotypes |  |  |  |  |  |  |  |  |  |  |  |  |
| --- | --- | --- | --- | --- | --- | --- | --- | --- | --- | --- | --- | --- |
| Location | Periplasm |  | Cytoplasm:<br>C-terminus |  | Cytoplasm:<br>Active Site |  |  |  |  |  |  |  |
| Mutation | Q395K |  | 526insA |  | G569C |  | G569V |  | G569D |  | P646S |  |
| Encoding<sup>1</sup> | chr | epi | chr | epi | chr | epi | chr | epi | chr | epi | chr | epi |
| Culture tackiness | + | + | + | = | + | + | + | + | + | + | + | + |
| Sedimentation efficiency<sup>2</sup> | + | - | - | = | - | - | - | - | - | = | - | - |
| Cell-free EPS | ++ | = | + | = | + | + | + | = | + | = | + | = |
| CPS | -- | + | - | = | = | + | - | = | - | = | - | = |
| Wzc-Y-P<sub>i</sub> | + | = | = | = | = | - | = | - | = | - | = | - |
| 616 mucoidy |  | - |  | - |  | + |  | + |  | + |  | + |
| Phenotype |  | part. |  | rec. |  | dom. |  | dom. |  | dom. |  | dom. |

<sup>1</sup>Chromosomal (chr) or episomal on pBAD18 (epi)
<sup>2</sup>Increased sedimentation efficiency (+) represents reduced mucoidy.
<sup>3</sup>Heterozygous phenotype in wildtype background: Partial penetrance (part.), recessive (rec.), or dominant (dom.)

Wzc_Q395K_ is the only periplasmic-localized point mutation identified in this study and behaves distinctly from the other identified Wzc mutations. The periplasmic domain is thought to coordinate CPS polymerization with secretion via the Wza pore.^21^ The phenotypes associated with the chromosomal versus episomal expression of the variant are distinct and informative. We speculate that the Wzc_Q395K_ phenotype exhibits partial penetrance when co-expressed with Wzc_WT_. Our model is that when only Wzc_Q395K_ is expressed in the cell, Wzc hyper-phosphorylation prevents re-assembly of the Wzc octamer and/or Wzc docking with Wza or Wzi. We suspect that the strong shift toward secretion of cell-free EPS may represent non-polymerized EPS sub-units secreted directly into the environment, which could explain the overt tackiness of these bacterial cultures. It is possible that when Wzc_Q395K_ is co-expressed with Wzc_WT_ that provides more balanced kinase-phosphatase activity, enabling CPS polymerization and anchoring to the cell surface. Overall, it seems that a minimal amount of cell-associated CPS is required for mucoidy and reduced sedimentation efficiency (**Fig. 5**), but that some other factor is also required. This remains in line with the current understanding of mucoidy in the field. We were unable to introduce targeted Wzc point mutations into the genome, which would be critical to test our model without the confounding effects of the transposon insertion sites. Our inability to generate these targeted mutations raises the question of whether the transposon insertions increase the frequency of Wzc mutation rate or temper any toxic effects of these mutations.

The bulk of the isolated Wzc point mutations are localized to the C-terminal active site in the cytoplasm. These include Wzc_G569C_, Wzc_G569V_, Wzc_G569D_, and Wzc_P646S_. Whether expressed as a secondary mutations on the chromosome or episomally in wildtype KPPR1, these mutations generally increased mucoidy and reduced Wzc-Y-P_i_ (**Table 2**). When they were expressed from the chromosome, cell-free EPS was increased, but this did not occur when expressed *in trans*. Furthermore, episomal expression of these active site mutations are sufficient to confer mucoidy to *K. variicola* strain 616, without significantly altering CPS or cell-free EPS production. These observations underscore that it is not cell-free EPS that drives mucoidy. One limitation to this interpretation is that CPS and EPS were quantified by detecting uronic acids with 2-hydroxydiphenyl. This assay is specific for CPS as acapsular strains have background levels of uronic acid;^18^ however, this does not preclude the possibility that sugars, not detected by 2-hydroxydiphenyl, drive the mucoid phenotype. The observation that expressing these four active site mutants *in trans* increases mucoidy and reduced Wzc-Y-P_i_ without impacting EPS and CPS, suggests that the Wzc tyrosine kinase may be a keystone regulator of *K. pneumoniae* mucoidy. The reduced Wzc-Y-P_i_ levels of these active site mutants reveals that these mutations drive Wzc to exist primarily in a de-phosphorylated state. Prior data support that Wzc dephosphorylation enables multimerization and that dynamic Wzc dissociation and multimerization is critical for CPS biosynthesis.^21^ These Wzc active site mutants may allow the protein to dephosphorylate more quickly, altering the export of cryptic non-capsular sugars or indirectly controling downstream tyrosine-phosphorylated targets that drive mucoidy. This model is in line with other evidence that mucoidy is regulated by post-translational modifications.^38^ For example, the minor but significant changes in *cps* gene expression (**Fig. 1C**) could be amplified by posttranslational phosphorylation. Nonetheless, we cannot discriminate whether the reduced Wzc-Y-P_i_ levels are due to quicker dephosphorylation or slower phosphorylation of Wzc.

The remaining Wzc mutation identified here inserts an alanine after residue 526 in the C-terminal cytoplasmic domain of the protein. This mutation appears to be recessive when expressed in a strain that also encodes wildtype Wzc. When the only copy of Wzc bears the 526_insA_ mutation, mucoidy and cell-free EPS increase (**Fig. 5A**), but these phenotypes are absent when Wzc_526insA_ is co-expressed in wildtype KPPR1, 616 or 1346. We confirmed that no other Wzc mutations are present in *pfkB-*Wzc_526insA_ or on the pBAD18-Wzc_526insA_ and that the protein is expressed from pBAD18 (**Fig. S2**). Despite exhibiting increased mucoidy, *pfkB-*Wzc_526insA_ does not impact Wzc-Y-P_i_ abundance (**Fig. 5A** and **7B**). This would suggest that Wzc phosphorylation status is not the only factor controlling mucoidy. It is possible that this mutation impacts Wzc multimerization or interactions with other binding partners. When co-expressed with Wzc_WT_ the associated phenotypes are lost, suggesting that this phenotype is recessive and can be compensated for by the presence of Wzc_WT_. A targeted Wzc_526insA_ mutant would validate if this model is correct, but as discussed above, we have been unable to introduce targeted Wzc point mutations into the chromosome.

Other studies have identified similar active site Wzc mutations that increase mucoidy in *K. pneumoniae* and *Acinetobacter baumannii*.^24,25,40^ Wzc mutations reported in *A. baumannii* increased mucoidy, increased production of higher molecular weight EPS, and was typically associated with reduced Wzc-Y-P_i_ abundance.^25^ However, some of the reported mutants did not reduce Wzc-Y-P_i_, akin what we have observed with Wzc_Q395K_ and Wzc_526insA_ in *K. pneumoniae* (**Fig. 7**). Furthermore, *A. baumannii* increased mucoidy and EPS production in response to chloramphenicol and erythromycin treatment, without impacting Wzc-Y-P_i_ abundance. Distinct from *K. pneumoniae*, no other tyrosine-phosphorylated proteins were detected in *A. baumannii*. Other studies have identified Wzc variants in *K. pneumoniae*, but none have examined the impact on cellular tyrosine phosphorylation.^24,40^ One study found that Wzc mutations increased mucoidy and total EPS in *rmp-*negative strains,^24^ while the other study found that Wzc mutations increased mucoidy, but not EPS, in both *rmp-*negative and - positive strains, including a *K. variicola* isolate.^40^ *K. pneumoniae* isolates with Wzc variants appear to be more resistant to phagocytosis and exhibit increased dissemination and virulence.^24^ Others have also noted the challenge of generating targeted Wzc point mutations, further strengthening our hypothesis that the transposon insertions may have provided some compensation for any lethal sideeffects of Wzc mutation.^24^ Alternatively, it could have been static culture in human urine that enriched for Wzc mutants, akin to the methods used in Nucci *et al*.^40^ Altogether, these data, along with ours, support that lower Wzc-Y-P_i_ abundance increases mucoidy in Gram-negative bacteria, but the biochemical mechanism driving mucoidy may differ in *K. pneumoniae* as lower Wzc-Y-P_i_ abundance does not always increase EPS production (**Fig. 6A-B**). It seems most likely that Wzc mutations that do not reduce Wzc-Y-P_i_ abundance, may increase mucoidy by altering Wzc protein-protein or protein-substrate interactions, which could be represented in the lower molecular weight bands observed by PY20 western blot. Previous studies have reported the tyrosine phosphorylation of other CPS biosynthesis proteins and found that abrogation of WcaJ phosphorylation reduced CPS production and virulence, so it is quite possible that tyrosine phosphorylation also impacts mucoidy, as some of our data suggest.^41^

In summary, our data have found that *K. pneumoniae* regulates mucoidy in response to environmental cues and implicate Wzc phosphosignaling and activity in controlling *K. pneumoniae* mucoidy. The active regulation of mucoidy in response to the environment raises questions regarding *K. pneumoniae* behavior during infection and whether mucoidy is actively suppressed during UTI, where it appears to be a disadvantage, or increased during dissemination.^24^ Furthermore, this active regulation of mucoidy suggests that discrete signals control the regulation of mucoidy, which may represent a potential anti-virulence target. It is possible that adventitious Wzc mutations drive the transition of *K. pneumoniae* to disseminate from the urinary tract, especially if the area is prone to mutagenesis.^24^ Finally, our data have shown that Wzc is sufficient to increase mucoidy independent of *rmpACD* and increased CPS biosynthesis. The sufficiency of Wzc to induce mucoidy independent of *rmpACD* suggests that Wzc functions downstream of RmpD in regulating mucoidy and that it may perform other cellular tasks besides regulating CPS biosynthesis. Therefore, Wzc activity may represent the lynchpin that coordinates both CPS biosynthesis and mucoidy.

## METHODS

### Bacterial strains and culture conditions

All primers, strains and plasmids described in these studies are detailed **S1** and **S2 Tables**. Bacteria were cultured in lysogeny broth (LB) (5 g/L yeast extract, 10 g/L tryptone, 0.5 g/L NaCl) at 200 rpm and 37°C, unless otherwise noted. Solid medium was prepared by the addition of 15 g/L bacto-agar prior to autoclaving. When appropriate, antibiotics were added at the following concentrations, kanamycin (25 μg/mL) and ampicillin (100 μg/mL). Human urine was collected anonymously from healthy women who were not menstruating, pregnant, nor within two weeks of antibiotic treatment. Urine was pooled from at least 5 independent donors and vacuum filtered and sterilized through 0.2 um PES membrane. Sterilized urine was stored in 50 mL aliquots at -20*C and working volumes were stored at 4*C. Solid urine medium was made by mixing warmed urine 1:1 with autoclaved bacto-agar (30 g/L) then poured into petri dishes.

### Uronic acid quantification

For experiments where bacteria were cultured in urine, 1 OD_600_ of bacteria was transferred to a microcentrifuge tube and pelleted at 21,000 *x g* for 15 min. The bacterial pellet was washed with 1 mL of PBS and centrifuged at 21,000 *x g* for 15 min. The washed cells were resuspended to 250 μL then 50 μL of 3-14 Zwittergent were added and uronic acid quantification was performed as previously described. For all other experiments, uronic acid quantification was performed as previously described.^18,42^ Cell-free EPS was quantified by mixing 250 μL of the overnight culture was mixed with 50 μL of ultra-pure water, instead of 3-14 Zwittergent. The culture was pelleted at 17,000 *x g* for 5 min then 100 μL of the upper supernatant was transferred to 400 μL of cold ethanol to precipitate cell-free EPS and uronic acid quantification was performed as previously described. CPS uronic acid content was deduced by subtracting the uronic acid levels in the whole culture from those in cell-free EPS.

### Hypermucoviscosity

Hypermucoviscosity was quantified as previously described (Mike PLOS Path). In brief, 1 OD_600_ unit of bacteria cultured overnight was pelleted in a 2 mL microcentrifuge tube at 1,000 *x g* for 5 min. The OD_600_ of the supernatant was then quantified. If experimental conditions resulted in an overnight OD_600_ <1 (e.g. urine), 1 OD_600_ of bacteria was transferred to a 2 mL microcentrifuge tube and pelleted at 21,000 *x g* for 15 min. All but 100 μL of the sample supernatant were removed, then the bacterial pellet was resuspended to 1 mL with PBS and sedimentation efficiency assessed as described above.

### RNA isolation and quantitative RT-PCR

qRT-PCR was performed as previously described with some modifications.^43^ In brief, bacteria cultured overnight in LB medium were sub-culture 1:100 into LB medium or 1:50 into sterile human urine. The bacteria were cultured with aeration at 37 °C, 200 RPM for 2 h then approximately 1×10^9^ CFU of bacteria were mixed at a 2:1 (vol:vol) ratio of RNAProtect (Qiagen) and incubated at room temperature for 5 min. Samples were then pelleted at 6,000 *x g* for 10 min and the supernatant was drained.

RNA was purified with the RNeasy mini-prep kit (Qiagen) after lysozyme and proteinase K treatment for 40 min at room temperature. Samples were subjected to on-column DNase I treatment at room temperature for 20 min according to the manufacturer’s directions, eluting twice with 30 μL of RNase-free water. cDNA synthesis was performed with SuperScript III Reverse Transcriptase (Invitrogen) on an equal amount of RNA for each sample, roughly 3 μg total. The resulting cDNA was diluted 1:50 in water and utilized for real-time quantitative reverse transcription PCR (qRT-PCR) in a QuantStudio 3 PCR system (Applied Biosystem) with SYBRGreen PowerUp reagent (Invitrogen), Primers used to amplify internal fragments of CPS biosynthesis genes are listed in **Table S1**. The *gap2* transcript was used as an internal control. The relative fold change was calculated using the comparative threshold cycle (*C*_*T*_) method.^44^ PCR amplification efficiency controls were performed for each primer set and dissociation curves were conducted to verify the amplification of a single product.

### Transposon screen in urine

The screen was performed as described previously, except cultures were grown in 100 μL of pooled and sterile-filtered human urine.^18^ The hits were struck onto urine agar, incubated at 37°C overnight and evaluated by string test the following day. At least three colonies of each primary hit were arrayed into a microplate to measure sedimentation efficiency in a secondary screen. Pathway analysis was performed using KEGG GENES (Kyoto Encyclopedia of Genes and Genomes).^45,46^

### Transposon site insertion validation

Bacterial lysates were used as genomic DNA template for PCR. Templates were prepared by inoculating 100 μL nuclease-free water with bacterial colonies, boiling the samples in the microwave for 1 min, then performing one freeze-thaw cycle. 1 μL of cell lysate was used as template for a 25 μL PCR reaction using *Taq* DNA Polymerase with ThermoPol Buffer (NEB). For each mutant, one primer annealed to the transposon sequence and one primer annealed to the genomic region adjacent to the predicted transposon insertion site (**Table S1**). PCR amplification was confirmed on a 1% agarose gel.

### Whole genome sequencing and analysis

Genomic DNA was purified from overnight cultures of each bacterial strain. 1 mL of bacterial culture was pelleted at 15,000 x *g* for 15 min at 4 °C then samples were prepared according to the directions for the Wizard HMW DNA Extraction kit for Gram-negative Bacteria (Promega). The University of Michigan Microbiome Core quantified the genomic DNA samples, prepared the libraries using an Illumina DNA prep kit, and sequenced with a MiSeq Reagent Kit v3 (600-cycle). Sequence variants were detected using the Variation Analysis pipeline on PATRIC with the *K. pneumoniae* subsp. *pneumoniae* strain VK055 [RefSeq Accession: NZ_CP009208.1; PATRIC Genome ID: 72407.38] as the target genome.^33,34^ Parameters used were the BWA-mem Aligner and FreeBayes SNP Caller.

### *wzc* Sanger sequencing and analysis

Cell lysates were prepared as described above used as DNA template for PCR amplification using Phusion polymerase (NEB) and oligonucleotides that anneal 150 bp outside of *wzc*. PCR amplification was confirmed on a 1% agarose gel, DNA fragments column purified (Epoch), and Sanger sequenced with nested primers. Point mutations were identified using ClustalW (MegAlign®. Version 15.3. DNASTAR. Madison, WI).

### Molecular cloning and transformation

Oligonucleotides and plasmids used in this study are listed in **Tables S1** and **S2**. A 2 kb fragment encompassing 1 kb upstream and 1 kb downstream of the *wzc* stop codon was TOPO-TA cloned into pCR2.1 (Invitrogen). Each Wzc point mutation was introduced either by inverse PCR with overlapping mutagenic primers and self-ligation or by inverse PCR with 5’-phosphorylated, non-overlapping primers and T4 ligation. The pBAD18 backbone, *wzc* 5’ fragment, and mutated *wzc* 3’ fragment were PCR amplified, gel purified (Monarch, NEB), and assembled NEBuilder HiFi DNA Assembly mix (NEB) for 1 h at 60 °C.^47^ The ligated products were dialyzed on 0.025 um MCE membranes (MilliporeSigma) against 10% glycerol prior to electroporation. A hexa-histidine tag was added to each pBAD18-Wzc vector with 5’-phosphorylated, non-overlapping primers and T4 ligation. The integrity of each plasmid was confirmed by sequencing.

Electroporation of vectors into TOP10 *E. coli* or *K. pneumoniae* complex strains were performed as previously described.^18^

### Genetic characterization of 616 and 1346

Raw sequence reads of UTI616 (SRA Accession: SRR4115176) and UTI1346 (SRA Accession: SRR4115182) from NCBI database were assembled using PATRIC (https://www.bv-brc.org). Assembled sequences were uploaded to *Pathogenwatch* (https://pathogen.watch/) for species identification.

### Western blot

Whole cell lystates were prepared from overnight cultures of each bacterial strain. One mL of bacterial culture was pelleted at 21,000 x *g* for 15 min at 4 °C and samples were kept on ice until they were boiled. Bacterial pellets were resuspended in 250 μL of lysis buffer; lysis buffer was prepared fresh as follows: one cOmplete Mini, EDTA-free Protease Inhibitor Cocktail tablet (Roche) and one PhosSTOP tablet (Roche) dissolvd in 7 mL BugBuster (MilliporeSigma). Each sample was sonicated in two, 5 sec pulses with 50% amplitude and 5 s rest between each pulse. Protein concentration was quantified by BCA assay (Pierce). Samples were prepared at 1 μg/μL in 1X SDS loading buffer and boiled at 70 °C for 3.5 min. Ten μg of protein were resolved on 12% SDS-PAGE gels and transferred to nitrocellulose. Membranes were blocked with 5% BSA in TBS overnight then washed with TBS-T and probed with 1:2,500 mouse anti-phosphotyrosine PY20 (primary) or 1:5,000 mouse anti-His-Tag (primary) then 1:5000 goat anti-mouse IgG(H+L)-HRP (secondary) (SouthernBiotech). Blots were developed with ECL Western Blotting Substrate (Pierce) and imaged on a G:Box Imager (Syngene).

### Western blot quantification

Western blots were quantified using ImageJ version 1.53K for Windows. In brief, an equal area of each band was measured and subtracted for respective background. Ratio of arbitrary value of each band to wild type was used in the quantification graph. Quantification shown in the graphs are average of triplicates.

### Statistics

All replicates represent biological replicates and were replicated at least three times. All statistical analyses were computed in Prism 5.03 (GraphPad Software, La Jolla, CA). For experiments comparing multiple groups, significance was calculated using two-way ANOVA with a Bonferroni post-test to compare specific groups. A one sample *t* test was applied when comparing two groups or a single group to a hypothetical value of 1.00. In all instances, results were considered significant if the P value was less than or equal to 0.05.

## ACKNOWLEDGEMENTS

The authors thank Dr. Michael Bachman for strains NTUH-K2044, 616 and 1346, and for bioinformatic guidance. The authors also thank members of the Mobley and Bachman labs at the University of Michigan and the Department of Medical Microbiology and Immunology at the University of Toledo for critical feedback and for sharing technical expertise and resources, especially Dr. Sébastien Crépin, Valerie Forsyth, Andrew Stark, Dr. Jyl Matson, and Dr. Robert Blumenthal. This research was supported by work performed by the University of Michigan Microbiome Core. Research reported in this publication was supported by K22 AI145849 to L.A.M. and R01 AI059722 to H.L.T.M from the National Institutes of Health. The content is solely the responsibility of the authors and does not necessarily represent the official views of the National Institutes of Health.

## Notes

### Competing Interest Statement

The authors have declared no competing interest.

