## Supplemental figures and tables for "Regulation of *Klebsiella pneumoniae* mucoidy by the bacterial tyrosine kinase Wzc"

### Supplementary Data

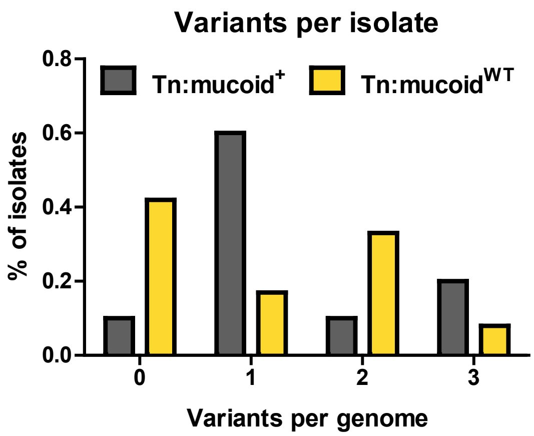


**Figure S1. Transposon isolates with elevated mucoidy have increased frequency of single genetic variations.** The genetic variation of 23 *K. pneumoniae* transposon isolates was determined using the Variation Analysis pipeline on PATRIC.[33,34] The number of non-synonymous mutations per genome was plotted versus the frequency that number of mutations occurred in Tn:mucoid$^{+}$ vs Tn:mucoid$^{WT}$ isolates.

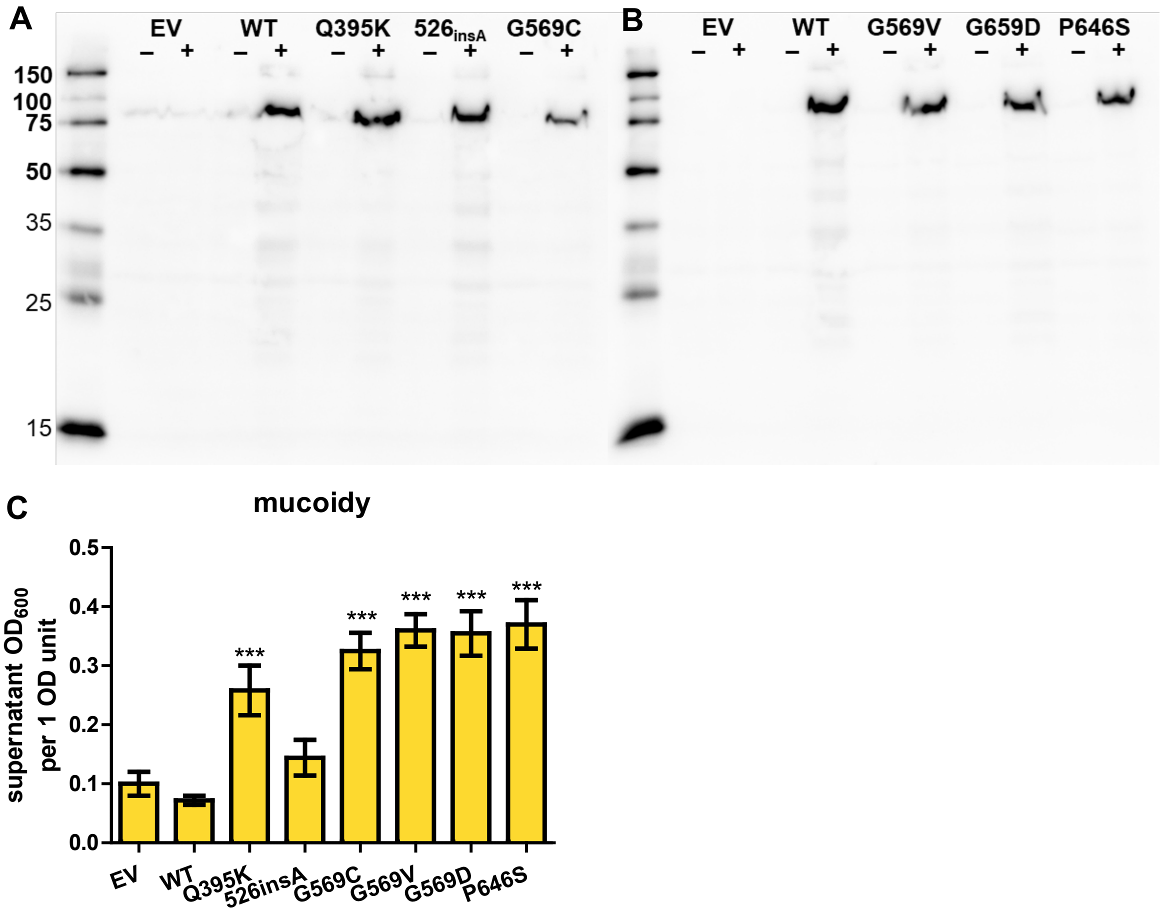


**Figure S2. Arabinose-induced expression of Wzc-His$_6$ tightly controls expression and mucoidy.** His$_6$-tagged wildtype (WT) Wzc and six variants (Q395K, 526$_{insA}$, G569C, G569V, G569D, P646S) were cloned under an arabinose-inducible promoter on pBAD18 then introduced into wildtype KPPR1. Empty vector pBAD18 (EV) was used as a negative control. (**A-B**) *K. pneumoniae* strains were cultured in LB medium containing kanamycin without (-) and with (+) 50 mM L-arabinose. Whole cell lysates were resolved by SDS-PAGE, transferred to nitrocellulose, then probed with anti-His$_6$ antibody. Lysates were prepared and analyzed ≥3 independent times. One representative image is shown. (**C**) The same strains were cultured in LB medium containing 50 mM L-arabinose and kanamycin. Mucoidy was determined by quantifying the supernatant $OD_{600}$ after sedimenting 1 $OD_{600}$ unit of culture at 1,000 *x g* for 5 min. Statistical significance was determined using two-way ANOVA with a Bonferroni post-test to compare specific groups. * $p < 0.05$; ** $p < 0.01$; *** $p < 0.001$; # $p < 0.0001$. Experiments were performed ≥3 independent times, in triplicate.

### Supplementary Tables

**S1 Table. Primers used in this study.**

| Primer Name | Sequence | Purpose | Source |
|---|---|---|---|
| LAM129 | TCGGCGACAACCCATTT | qPCR_VK055_5012_galF_F | This study |
| LAM130 | GATCATAGCGGCGAGGTTATAG | qPCR_VK055_5012_galF_R | This study |
| LAM175 | TGAACTGGCAACTCATCTCAT | qPCR_VK055_5013_F2 | This study |
| LAM176 | CGCAGCATCAGGACAATAAAC | qPCR_VK055_5013_R2 | This study |
| LAM133 | TAGGTGCATCGCGCATTAT | qPCR_VK055_5014_wzi_F | This study |
| LAM134 | GTTCGTTATCTGTCCCGGTATT | qPCR_VK055_5014_wzi_R | This study |
| LAM135 | AATGGACCGCAGTGGTATG | qPCR_VK055_5015_wza_F | This study |
| LAM136 | CACCTTTCAACTGGCGAATAAC | qPCR_VK055_5015_wza_R | This study |
| LAM137 | GGATTTCATTGGCAGGTCATTT | qPCR_VK055_5016_wzb_F | This study |
| LAM138 | CGGGTGCAATGTTGCTTATC | qPCR_VK055_5016_wzb_R | This study |
| LAM139 | CGTGATGTTGAGTCAGGTAGAG | qPCR_VK055_5017_etk_F | This study |
| LAM140 | TGGGTTACAGCGCTATCAATTA | qPCR_VK055_5017_etk_R | This study |
| LAM141 | TCATCCATATCGCGGAAACC | qPCR_VK055_5018_F | This study |
| LAM142 | TTGATTGACTCTCAGGGCATATT | qPCR_VK055_5018_R | This study |
| LAM143 | TTTATCGGGCGGTTGACTTAC | qPCR_VK055_5019_F | This study |
| LAM144 | TCAAACTCTCCTGCAGCAATAA | qPCR_VK055_5019_R | This study |
| LAM145 | CCAACCATGGTGGTCGATTA | qPCR_VK055_5020_F | This study |
| LAM146 | GGCTTCGTCTATTTCCTCCTTAT | qPCR_VK055_5020_R | This study |
| LAM147 | GCTGTCGCTATGCTCAATAGTA | qPCR_VK055_5021_F | This study |
| LAM148 | CTTGTTGCCACGGATGAATATG | qPCR_VK055_5021_R | This study |
| LAM177 | CTTCATGGATGAGGCTGGATAG | qPCR_VK055_5022_wzx_F2 | This study |
| LAM178 | CTGGCCATGAGGCATGATAATA | qPCR_VK055_5022_wzx_R2 | This study |
| LAM179 | TCGGGAGGAATGATGTTGAAG | qPCR_VK055_5023_F2 | This study |
| LAM180 | TAGTCAGAGGCAGAACAATTCC | qPCR_VK055_5023_R2 | This study |
| LAM153 | GGGTTGACATTGCTGACTCTA | qPCR_VK055_5024_F | This study |
| LAM154 | GAACCCTCCAAGAAACCATACT | qPCR_VK055_5024_R | This study |
| LAM155 | TGGCGTGGTGAAACAGATAC | qPCR_VK055_5025_wcaJ_F | This study |
| LAM156 | CCAAATGCTCCAACCTCTGATA | qPCR_VK055_5025_wcaJ_R | This study |
| LAM157 | GCAGCTACCTGATCGACATTAC | qPCR_VK055_5026_gnd_F | This study |
| LAM158 | CCTTTGTTTGCCGCTTCATC | qPCR_VK055_5026_gnd_R | This study |
| LAM159 | CCGCCGACCATATCATCAATA | qPCR_VK055_5027_manC_F | This study |
| LAM160 | GATACCGAAGGTCACCAGATG | qPCR_VK055_5027_manC_R | This study |
| LAM161 | GCGGCACGGAAGAGATTTA | qPCR_VK055_5028_manB_F | This study |
| LAM162 | CCGTTGTAGTTCATCGGGTTAT | qPCR_VK055_5028_manB_R | This study |
| LAM163 | TACGGTGGAAGCGGTTATTC | qPCR_VK055_5029_ugd_F | This study |
| LAM164 | TCTTTGATGTCGCGGGTAAA | qPCR_VK055_5029_ugd_R | This study |
| LAM185 | CGCTGCTAAAGCAGTAGGTAAA | qPCR_VK055_1263_gap2_F | This study |
| LAM186 | CGGTCAGGTCAACAACAGATAC | qPCR_VK055_1263_gap2_R | This study |

**S1 Table cont'd. Primers used in this study.**

| Primer Name | Sequence | Purpose | Source |
|---|---|---|---|
| LAM187 | TCTTCGTATGCCGTCTTCTGCTTG | Tn verification, anneals to Tn | [18] |
| LAM228 | GCAATTCGTCGTATAAGAAAGAATGCGAAGACG | Tn verification, anneals 100 bp upstream of pheA start | This study |
| LAM214 | GACCTTATCATGCGCGACGG | Tn verification, anneals 260 bp upstream of gmhA2 start | This study |
| LAM215 | GCACGAGGGCCAGACG | Tn verification, anneals 280 bp downstream of rafA start | This study |
| LAM216 | CCGCACTCATATGCGGAGAAATAAATTACTGC | Tn verification, anneals 145 bp upstream of VK055_2735 sta | This study |
| LAM217 | GTTGTTGCTCCACAGGCCAG | Tn verification, anneals 75 bp downstream of VK055_1828 s | This study |
| LAM218 | GCTCAGGGCTTTTTTGTTTGATCTGTTCTC | Tn verification, anneals 150 bp upstream of VK055_2345 sta | This study |
| LAM219 | CCAAAGATGCGATAACAACCCATGAACAG | Tn verification, anneals 460 bp upstream of VK055_4105 sta | This study |
| LAM220 | CCCTAGAACCGGCGAGC | Tn verification, anneals 180 bp upstream of VK055_0204 sta | This study |
| LAM221 | CTTCCACCTGCAGACCGTTAGG | Tn verification, anneals 360 bp upstream of dtpD start | This study |
| LAM222 | GAGAGGAGCAAGTCGACCACC | Tn verification, anneals 440 bp upstream of VK055_4261 sta | This study |
| LAM223 | GTGCAACCCCAGCTTAATCACC | Tn verification, anneals 280 bp upstream of VK055_0874 sta | This study |
| LAM224 | CACCGTCATCTCCACCTTTGGTC | Tn verification, anneals 275 bp upstream of nouN start | This study |
| LAM225 | GTCACCGTCTGCGTTTTACCG | Tn verification, anneals 100 bp upstream of pfkB start | This study |
| LAM226 | CTTCCGCTGCCTGCGTC | Tn verification, anneals 360 bp upstream of proB start | This study |
| LAM227 | GTAGATGGATTGTGGGATTTAGCGAAAGTAGC | Tn verification, anneals 46 bp downstream of VK055_1417 s | This study |
| LAM229 | GTTGTTTGGCCATTGGTTAG | *wzc* sequencing, anneals 150 bp upstream of *wzc* start | This study |
| LAM230 | GACTATCAATTTCATTGTTTTCCG | *wzc* sequencing, anneals 150 bp downstream of *wzc* stop | This study |
| LAM231 | GTGAAGAGGCCTTTGCATC | *wzc* sequencing, anneals 100 bp upstream of *wzc* start | This study |
| LAM232 | ATCATATCTAGGCGATGATCC | *wzc* sequencing, anneals 700 bp downstream of *wzc* start | This study |
| LAM233 | AACGAGATGACAGTAGACCTAAGGT | *wzc* sequencing, anneals 1450 bp downstream of *wzc* star | This study |
| LAM388 | CACATAGCTATTTACCATATCGCTTTTCC | Rev primer anneals 1 kb downstream of *wzc* stop | This study |
| LAM389 | AAGGAAAACATTACAGGATGAAAGAGG | Fwd primer anneals 1 kb upstream of *wzc* stop | This study |
| LAM394 | GCAATTATTAAATCGACAGAAAGAATTGAATATTGC | wzc_Q395K_Fwd for inverse PCR | This study |
| LAM395 | GCAATATTCAATTCTTTCTGTCGATTTAATAATTGC | wzc_Q395K_Rev for inverse PCR | This study |
| LAM436 | GCAGCAAGAAATAATCTCTTAATGATTTCCG | wzc_526insA_Fwd phosphorylated primer for inverse PCR | This study |
| LAM437 | TTCAAGCATTGCAAAGTGCAG | wzc_526insA_Rev phosphorylated primer for inverse PCR | This study |
| LAM398 | CGATGCTGATTTAAGGAAATGTTATACTCAC | wzc_G569C_Fwd for inverse PCR | This study |
| LAM399 | GTGAGTATAACATTTCCTTAAATCAGCATCG | wzc_G569C_Rev for inverse PCR | This study |
| LAM434 | TTTATACTCACAAATTATTTAATATAAAAAACAC | wzc_G569V_Fwd phosphorylated primer for inverse PCR | This study |
| LAM435 | CTTTCCTTAAATCAGCATCGATAAAAATAAC | wzc_G569V_Rev phosphorylated primer for inverse PCR | This study |
| LAM400 | CGATGCTGATTTAAGGAAAGATTATACTCAC | wzc_G569D_Fwd for inverse PCR | This study |
| LAM401 | GTGAGTATAATCTTTCCTTAAATCAGCATCG | wzc_G569D_Rev for inverse PCR | This study |
| LAM404 | CATATTAGATACACCTTCAATCCTGGCTGTG | wzc_P646S_Fwd for inverse PCR | This study |
| LAM405 | CACAGCCAGGATTGAAGGTGTATCTAATATG | wzc_P646S_Rev for inverse PCR | This study |
| SDH_P17 | CAGCCTGATACAGATTAAAT | pBAD18Kan_fwd for NEBuilder vector | [47] |
| SDH_P18 | GGAGAAACAGTAGAGAGTTG | pBAD18Kan_rev for NEBuilder vector | [47] |
| LAM441 | caactctctactgtttctccATGACTTCAATATCCAAAAAGAAG | wzc_5'_fwd for NEBuilder insert fragment 1 | This study |
| LAM442 | ggcattgttgCAACACGCTTATTTAATTTCC | wzc_5'_rev for NEBuilder insert fragment 1 | This study |
| LAM443 | aagcgtgttgCAACAATGCCTGAGACTC | wzc_3'_fwd for NEBuilder insert fragment 2 | This study |
| LAM444 | atttaatctgtatcaggctgCTATTTTTTATCTGAATATGAATAATCG | wzc_3'_rev for NEBuilder insert fragment 2 | This study |
| LK19 | CATCACCATCACCATCACTAGCAGCCTGATACAGATTAAATC | pBAD18-CtermHis6_Fwd, 5' phosphorylated | This study |
| LK20 | TTTTTTATCTGAATATGAATAATCGTAATAATTATATCC | WzcCterm_Rev, 5' phosphorylated | This study |

| S2 Table. Strains and plasmids used in this study | | | | | | | | |
|---|---|---|---|---|---|---|---|---|
| Strain | Plasmid | Relevant details[a] | Source | | Strain | Plasmid | Relevant details[a] | Source |
| *Escherichia coli* | | | | | *Klebsiella pneumoniae* | | | |
| TOP10 | pCR2.1-Wzc$^{3'}$-$^{5'}$VK055_5018 | Amp$^r$ | This study | | KPPR1 | | ATCC 43816, Rif$^r$ | [27] |
| TOP10 | pCR2.1-Wzc$^{3'}_{Q395K}$-$^{5'}$VK055_5018 | Amp$^r$ | This study | | NTUH-K2044 | | | [28] |
| TOP10 | pCR2.1-Wzc$^{3'}_{526_insA}$-$^{5'}$VK055_5018 | Amp$^r$ | This study | | 616 | | Amp$^r$ | [26] |
| TOP10 | pCR2.1-Wzc$^{3'}_{G569C}$-$^{5'}$VK055_5018 | Amp$^r$ | This study | | 1346 | | Amp$^r$ | [26] |
| TOP10 | pCR2.1-Wzc$^{3'}_{G569V}$-$^{5'}$VK055_5018 | Amp$^r$ | This study | | KPPR1 | pBAD18 | Km$^r$, Rif$^r$ | This study |
| TOP10 | pCR2.1-Wzc$^{3'}_{G569D}$-$^{5'}$VK055_5018 | Amp$^r$ | This study | | KPPR1 | pBAD18-Wzc | Km$^r$, Rif$^r$ | This study |
| TOP10 | pCR2.1-Wzc$^{3'}_{P646S}$-$^{5'}$VK055_5018 | Amp$^r$ | This study | | KPPR1 | pBAD18-Wzc$_{Q395K}$ | Km$^r$, Rif$^r$ | This study |
| TOP10 | pBAD18-Wzc | Km$^r$ | This study | | KPPR1 | pBAD18-Wzc$_{526_insA}$ | Km$^r$, Rif$^r$ | This study |
| TOP10 | pBAD18-Wzc$_{Q395K}$ | Km$^r$ | This study | | KPPR1 | pBAD18-Wzc$_{G569C}$ | Km$^r$, Rif$^r$ | This study |
| TOP10 | pBAD18-Wzc$_{526_insA}$ | Km$^r$ | This study | | KPPR1 | pBAD18-Wzc$_{G569V}$ | Km$^r$, Rif$^r$ | This study |
| TOP10 | pBAD18-Wzc$_{G569C}$ | Km$^r$ | This study | | KPPR1 | pBAD18-Wzc$_{G569D}$ | Km$^r$, Rif$^r$ | This study |
| TOP10 | pBAD18-Wzc$_{G569V}$ | Km$^r$ | This study | | KPPR1 | pBAD18-Wzc$_{P646S}$ | Km$^r$, Rif$^r$ | This study |
| TOP10 | pBAD18-Wzc$_{G569D}$ | Km$^r$ | This study | | KPPR1 | pBAD18-Wzc-His$_6$ | Km$^r$, Rif$^r$ | This study |
| TOP10 | pBAD18-Wzc$_{P646S}$ | Km$^r$ | This study | | KPPR1 | pBAD18-Wzc$_{Q395K}$-His6 | Km$^r$, Rif$^r$ | This study |
| TOP10 | pBAD18-Wzc-His$_6$ | Km$^r$ | This study | | KPPR1 | pBAD18-Wzc$_{526_insA}$-His6 | Km$^r$, Rif$^r$ | This study |
| TOP10 | pBAD18-Wzc$_{Q395K}$-His6 | Km$^r$ | This study | | KPPR1 | pBAD18-Wzc$_{G569C}$-His6 | Km$^r$, Rif$^r$ | This study |
| TOP10 | pBAD18-Wzc$_{526_insA}$-His6 | Km$^r$ | This study | | KPPR1 | pBAD18-Wzc$_{G569V}$-His6 | Km$^r$, Rif$^r$ | This study |
| TOP10 | pBAD18-Wzc$_{G569C}$-His6 | Km$^r$ | This study | | KPPR1 | pBAD18-Wzc$_{G569D}$-His6 | Km$^r$, Rif$^r$ | This study |
| TOP10 | pBAD18-Wzc$_{G569V}$-His6 | Km$^r$ | This study | | KPPR1 | pBAD18-Wzc$_{P646S}$-His6 | Km$^r$, Rif$^r$ | This study |
| TOP10 | pBAD18-Wzc$_{G569D}$-His6 | Km$^r$ | This study | | 616 | pBAD18 | Km$^r$, Amp$^r$ | This study |
| TOP10 | pBAD18-Wzc$_{P646S}$-His6 | Km$^r$ | This study | | 616 | pBAD18-Wzc | Km$^r$, Amp$^r$ | This study |
| | | | | | 616 | pBAD18-Wzc$_{Q395K}$ | Km$^r$, Amp$^r$ | This study |
| | | | | | 616 | pBAD18-Wzc$_{526_insA}$ | Km$^r$, Amp$^r$ | This study |
| | | | | | 616 | pBAD18-Wzc$_{G569C}$ | Km$^r$, Amp$^r$ | This study |
| | | | | | 616 | pBAD18-Wzc$_{G569V}$ | Km$^r$, Amp$^r$ | This study |
| | | | | | 616 | pBAD18-Wzc$_{G569D}$ | Km$^r$, Amp$^r$ | This study |
| | | | | | 616 | pBAD18-Wzc$_{P646S}$ | Km$^r$, Amp$^r$ | This study |
| | | | | | 1346 | pBAD18 | Km$^r$, Amp$^r$ | This study |
| | | | | | 1346 | pBAD18-Wzc | Km$^r$, Amp$^r$ | This study |
| | | | | | 1346 | pBAD18-Wzc$_{Q395K}$ | Km$^r$, Amp$^r$ | This study |
| | | | | | 1346 | pBAD18-Wzc$_{526_insA}$ | Km$^r$, Amp$^r$ | This study |
| | | | | | 1346 | pBAD18-Wzc$_{G569C}$ | Km$^r$, Amp$^r$ | This study |
| | | | | | 1346 | pBAD18-Wzc$_{G569V}$ | Km$^r$, Amp$^r$ | This study |
| | | | | | 1346 | pBAD18-Wzc$_{G569D}$ | Km$^r$, Amp$^r$ | This study |
| | | | | | 1346 | pBAD18-Wzc$_{P646S}$ | Km$^r$, Amp$^r$ | This study |

[a]Km = kanamycin; Amp = ampicillin; Rif = rifampin
